# USP37 prevents premature disassembly of stressed replisomes by TRAIP

**DOI:** 10.1101/2024.09.03.611025

**Authors:** Olga V. Kochenova, Giuseppina D’Alessandro, Domenic Pilger, Ernst Schmid, Sean L. Richards, Marcos Rios Garcia, Satpal S. Jhujh, Andrea Voigt, Vipul Gupta, Christopher J. Carnie, R. Alex Wu, Nadia Gueorguieva, Grant S. Stewart, Johannes C. Walter, Stephen P. Jackson

**Author notes:** Equal contribution (Olga V. Kochenova, Giuseppina D’Alessandro, Domenic Pilger, Ernst Schmid).

## Abstract

The E3 ubiquitin ligase TRAIP associates with the replisome and helps this molecular machine deal with replication stress. Thus, TRAIP promotes DNA inter-strand crosslink repair by triggering the disassembly of CDC45-MCM2-7-GINS (CMG) helicases that have converged on these lesions. However, disassembly of single CMGs that have stalled temporarily would be deleterious, suggesting that TRAIP must be carefully regulated. Here, we demonstrate that human cells lacking the de-ubiquitylating enzyme USP37 are hypersensitive to topoisomerase poisons and other replication stress-inducing agents. We further show that TRAIP loss rescues the hypersensitivity of *USP37* knockout cells to topoisomerase inhibitors. In *Xenopus* egg extracts depleted of USP37, TRAIP promotes premature CMG ubiquitylation and disassembly when converging replisomes stall. Finally, guided by AlphaFold-Multimer, we discovered that binding to CDC45 mediates USP37’s response to topological stress. In conclusion, we propose that USP37 protects genome stability by preventing TRAIP-dependent CMG unloading when replication stress impedes timely termination.

## Main

Faithful DNA replication is essential to maintain genome integrity and prevent cancer and other diseases. Cells prepare for DNA replication in the G1 phase of the cell cycle, when ORC, CDC6 and CDT1 recruit MCM2-7 double hexamers to origins of DNA replication (licensing). In S phase, cyclin-dependent kinases (CDKs) and DBF4-dependent kinases (DDKs) cooperate with additional factors to recruit the tetrameric GINS complex (SLD5, PSF1, PSF2, and PSF3) and CDC45 to MCM2-7, leading to the formation and activation of two CMG helicases. Active CMGs translocate in the 3’ to 5’ direction along the leading-strand DNA template. Once CMG unwinds the replication origin, replisomes are assembled, and bi-directional DNA synthesis commences. Replication finishes when replisomes from adjacent origins converge, leading to termination^1–3^.

Replication termination is a highly regulated process that is critical for accurate genome maintenance^4^. To initiate termination, forks must merge, a process that depends critically on topoisomerase activity^5,6^. When converging CMGs meet, they pass each other, leading to disengagement of the lagging strand template from the outer face of CMG. Strand disengagement allows binding of the E3 ubiquitin ligase CRL2^Lrr^^1^ to CMG, leading to MCM7 ubiquitylation and CMG extraction from chromatin by the p97 ATPase (aka VCP)^7–9^. In the presence of DNA damage, CMGs can also dissociate from chromatin. For example, if CMG encounters a nick in the leading strand template, it slides off the end of the DNA, and if the nick is in the lagging strand template, lagging strand disengagement from CMG promotes its unloading by CRL2^Lrr^^1^ (Ref.^10^). Thus, while CRL2^Lrr1^ removes the bulk of CMGs from chromatin during a normal S phase, it also removes CMG at specific types of DNA damage.

CMG unloading is also promoted by a second RING E3 ubiquitin ligase called TRAIP. TRAIP associates with replisomes and ubiquitylates CMGs that have converged on a DNA inter-strand cross-link (ICL), triggering their unloading by p97 and activation of ICL repair (Extended Data Fig. 1ai and Ref.^11^). TRAIP also ubiquitylates DNA protein cross-links (DPCs) that block replisome progression (Extended Data Fig. 1aii), and it might also act on RNAPII complexes during transcription-replication conflicts^12,13^. Together, these observations suggest that in S phase, TRAIP ubiquitylates any proteinaceous structure encountered by the replisome (“*trans* ubiquitylation”)^14^. At the same time, TRAIP appears to be unable to ubiquitylate the replisome with which it travels (“*cis* ubiquitylation”; Extended Data Fig. 1aiii and Ref.^12^). This restriction of *cis* ubiquitylation is crucial to prevent premature CMG disassembly in S phase, which would lead to fork collapse and incomplete DNA replication. Indeed, in most instances of replication stress, which are ultimately overcome, CMG is likely not unloaded so that replication can resume after the stress is resolved^15–18^. Notably, once cells enter mitosis, TRAIP supports *cis* ubiquitylation of CMGs, which disengages remaining replisomes to allow orderly chromosome segregation (Extended Data Fig. 1b; Ref.^19–22)^. While TRAIP has the capacity for various forms of replisome-associated ubiquitylation, it remains unclear how aberrant TRAIP-mediated CMG ubiquitylation in interphase is avoided.

E3 ubiquitin ligases are counteracted by a family of ∼100 deubiquitylases (DUBs) – specialized proteases that can cleave isopeptide bonds between ubiquitin and lysine residues, which removes or remodels ubiquitin chains^23^. Ubiquitin-specific proteases (USPs) comprise the largest subfamily of DUBs, and their aberrant expression has been linked to the occurrence and progression of cancer, rendering them promising targets for anticancer therapies^24,25^. In particular, the DUB USP37 has been linked to the DNA damage response (DDR), genome stability and DNA replication^26–29^, and its overexpression provides a survival advantage to cancer cells^30–35^. Additional evidence points to USP37 as a possible replisome component^7,36,37^, yet its specific role(s) at the replisome remain unclear.

Here, we report that USP37 protects cells from replication stress and that TRAIP loss rescues the hypersensitivity of *USP37* knockout cells to inhibitors of topoisomerases. Using *Xenopus* egg extracts, we demonstrate that TRAIP ubiquitylates CMGs when replication forks stall at DPCs and/or experience topological stress, and that USP37 counteracts TRAIP-mediated CMG-ubiquitylation. Furthermore, using *in silico* screening for protein-protein interactions and site-directed mutagenesis, we provide evidence that USP37 mediates its function at the replisome by binding to CDC45. Based on these observations, we propose that USP37 promotes genome integrity and cell survival by preventing premature disassembly of stressed replisome by TRAIP.

## Results

### CRISPR screening identifies USP37 as promoting camptothecin resistance

To identify novel regulators of DNA topological stress, we performed a genome-scale CRISPR-Cas9 gene-inactivation cell fitness screen^38^ in human U2OS cells exposed to the TOP1 inhibitor camptothecin (CPT; Fig. 1a and Extended Data Fig. 2a). As expected, we identified factors involved in single-strand DNA break repair whose inactivation is known to cause CPT hypersensitivity, such as TDP1, PARP1 and XRCC1 (Fig. 1a). This analysis also identified the DUB USP37. To extend these results, we used CRISPR-Cas9 genome engineering to inactivate *USP37* in *TP53*-null human RPE-1 cells (Extended Data Fig. 2b) and observed that, compared to *USP37*-proficient control cells (CTRL), two independent *USP37* knockout clones (10 and 19) were hypersensitive to low doses of CPT (Fig. 1b). Additionally, these *USP37* knockout cells were hypersensitive to the TOP2 inhibitors etoposide and ICRF-193 (Fig. 1c-d), as well as the replicative DNA polymerase inhibitor aphidicolin (Fig. 1e). We further validated our observations in U2OS cells, where USP37 loss conferred hypersensitivity towards CPT, etoposide, aphidicolin and hydroxyurea, which impairs DNA replication by depleting dNTPs, but not towards the PARP1/2 inhibitor talazoparib, which causes replication-associated DNA double-strand breaks (Extended Data Fig. 2c-h). Collectively, these results suggested that USP37 generally mitigates the deleterious effects of replication fork stalling, but not necessarily fork breakage.

**Fig. 1:**
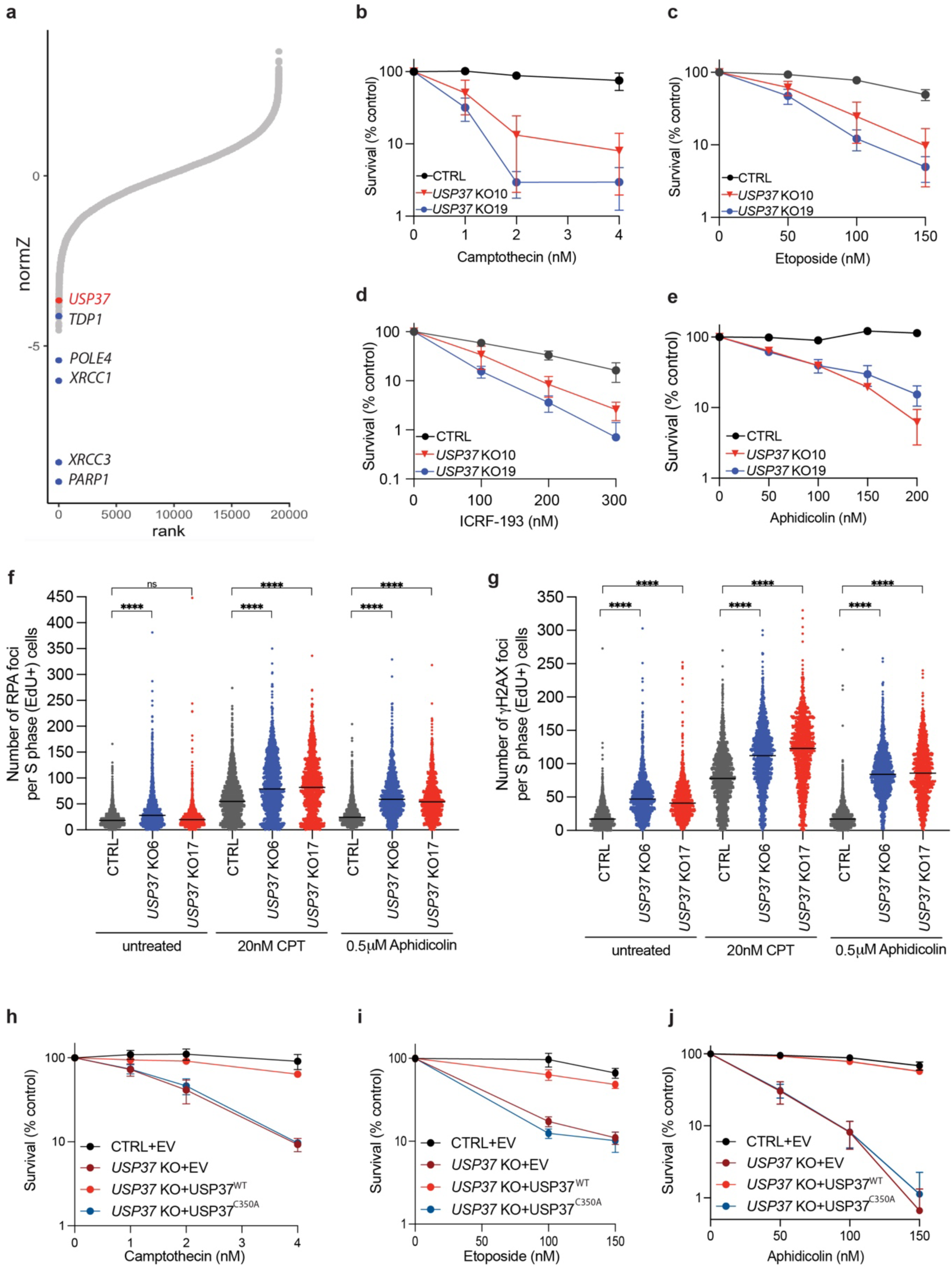
USP37 loss protects cells from topoisomerase poisons- and replication stress-associated DNA damage. **a,** Rank-plot showing gene enrichment scores (normZ) of CRISPR screen hits upon camptothecin treatment in human U2OS cells. Negative scores represent dropouts; that is genes whose loss is predicted to promote drug sensitivity. Blue dots indicate well characterised factors affecting cellular sensitivity to camptothecin; the red dot indicates *USP37*. **b-e,** Clonogenic survival assays of control (CTRL) and *USP37* knockout (KO; results from two independent clones plotted) RPE-1 *TP53* KO cells upon treatment with (**b**) camptothecin, (**c**) etoposide, (**d**) ICRF-193, or (**e**) aphidicolin. n=3 independent experiments. **f-g,** Quantification of RPA (**f**) or γH2AX (**g**) foci in S-phase cells (EdU positive). Cells were treated with the indicated doses of the drugs for 4h. n ≥ 4 independent experiments. Bars represent median. CPT is camptothecin. **h-j,** Clonogenic survival assays of control (CTRL) cells or *USP37* KO (clone 10) cells complemented with vectors expressing mCherry (EV), mCherry-USP37^WT^ or mCherry-USP37^C350A^ (catalytic inactive) upon treatment with camptothecin (**h**), etoposide (**i**), or aphidicolin (**j**); n=3 independent experiments. Bars represent means ± SEM.

To monitor the impact of USP37 loss on genome stability, we tested the levels of RPA and γH2AX foci, markers of ssDNA and DSB, respectively, in S-phase CTRL or *USP37* KO cells upon treatment with CPT or aphidicolin (Fig. 1f-g). Upon treatment with CPT, *USP37* KO cells showed increased RPA and γH2AX foci relative to CTRL cells. Similarly, treatment with aphidicolin, at doses that did not induce any damage in CTRL cells, strongly increased the number of RPA and γH2AX foci in *USP37* KO cells, in line with published data^29^. Overall, these data support a role for USP37 in promoting genome integrity upon treatment with CPT and aphidicolin.

To address whether the catalytic activity of USP37 is necessary to modulate its cellular functions, we generated a catalytically inactive point mutant (USP37^C350A^), as previously described^37^, and expressed it in *USP37* knockout cells alongside the wild-type USP37 (USP37^WT^) or an mCherry expressing vector (EV) (Extended Data Fig. 2i). We observed that overexpression of USP37^WT^ restored the viability of the knockout cells upon treatment with CPT (Fig. 1h), etoposide (Fig. 1i), or aphidicolin (Fig. 1j), while overexpression of the inactive mutant did not. These data indicated that the catalytic activity of USP37 protects cells from the deleterious effects of replication fork stalling.

### Genetic screening unveils functional connections between USP37 and TRAIP

To identify factors that functionally interact with USP37, we performed parallel genome-scale CRISPR-Cas9 knockout screens in wild-type (WT) and *USP37* knockout cells and looked for genes whose loss affected fitness differentially in the two genetic backgrounds. We found that TRAIP loss causes a fitness defect in WT but not *USP37* knockout U2OS cells (Fig. 2a). We validated this observation by performing cell-growth competition assays in Cas9-expressing WT and *USP37* knockout cells transiently transfected with a CRISPR single-guide RNA (sgRNA) targeting *TRAIP*. In line with previous reports^13,39,40^, *TRAIP* inactivation was not well tolerated in WT cells, as indicated by reduction over time of the cell population harboring mutations at the targeted *TRAIP* locus (“edited”). By contrast, such a reduction in the *TRAIP* edited population was not evident in the *USP37* knockout cells (Fig. 2b and Extended Data Fig. 3a). This epistatic relationship between *TRAIP* and *USP37* suggested that TRAIP and USP37 control a common pathway.

**Fig. 2:**
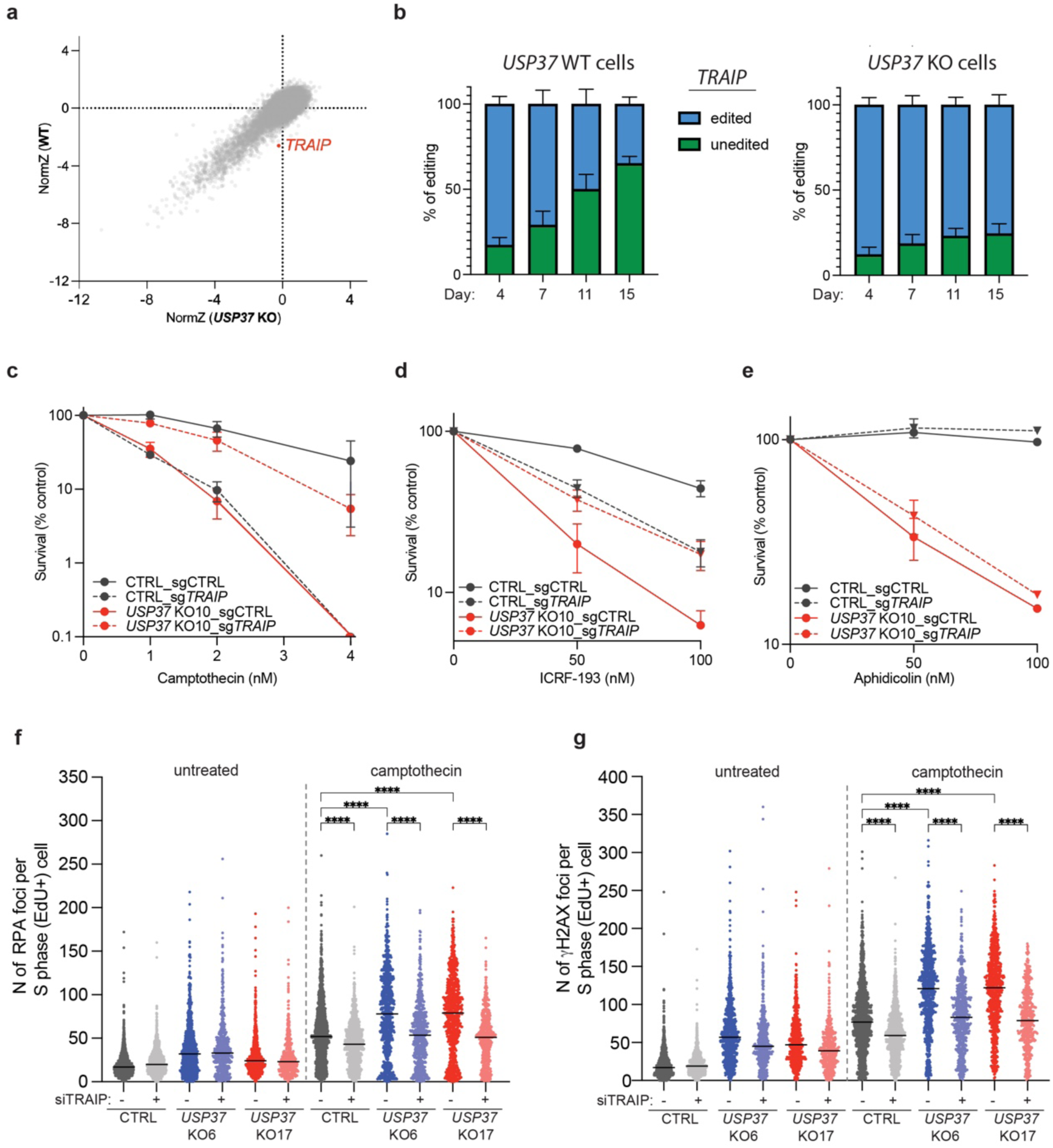
Genetic screen unveils functional connections between USP37 and TRAIP. **a,** Biplot showing gene enrichment scores (normZ) reflecting viability in untreated conditions of WT (y-axis) and *USP37* KO (x-axis) U2OS cells. **b,** U2OS *USP37* WT cells (left histogram) and *USP37* KO cells (right histogram) were transfected with a CRISPR sgRNA targeting *TRAIP* and samples were collected 4, 7, 11 and 15 days afterwards and subjected to *TRAIP* sequence analysis. Percentages of unedited or edited cells at the *TRAIP* locus at the indicated timepoints are plotted. **c-e,** Clonogenic survival assays of CTRL or *USP37* KO RPE-1 *TP53* KO cells transduced with a LacZ control sgRNA or with a sgRNA targeting *TRAIP* upon treatment with camptothecin (**c**), ICRF-193 (**d**), and aphidicolin (**e**). n=3 independent experiments. Bars represent means ± SEM. Half plot points indicate zero percent viability. **f-g,** Quantification of RPA (**f**) or γH2AX (**g**) foci in S-phase cells (EdU positive). Cells were treated with 20nM camptothecin for 4h. n = 3 independent experiments. Bars represent median.

Considering the above data, we tested whether TRAIP loss might affect the hypersensitivity of *USP37* knockout cells to CPT. To this end, we generated polyclonal knockouts of *TRAIP* in WT and *USP37* knockout *TP53*-null human RPE-1 cell backgrounds (Extended Data Fig. 3b) and subjected them to clonogenic survival assays. While *TRAIP* knockout and *USP37* knockout cells were hypersensitive to CPT, as expected^27,41^, the *USP37-TRAIP* double knockout cells largely lost their hypersensitivity towards CPT (Fig. 2c and Extended Data Fig. 4a), indicating that TRAIP loss compensates for the absence of USP37 and vice versa. Furthermore, while loss of USP37 or TRAIP alone caused increased sensitivity to ICRF-193, cells lacking both factors were more resistant than *USP37* knockout cells, and equally sensitive to the *TRAIP* knockout alone (Fig. 2d and Extended Data Fig. 4b). These findings therefore indicated that the main source of toxicity in *USP37* knockout cells upon treatment with CPT or ICRF-193 is due to TRAIP. In contrast, *TRAIP* loss did not restore *USP37* knockout cells’ tolerance of aphidicolin (Fig. 2e and Extended Data Fig. 4c). TRAIP depletion significantly reduced the number of CPT-induced RPA and γH2AX foci in CTRL cells (Fig. 2f-g and Extended Data Fig. 4d), consistent with previous findings^42^. This reduction was even more pronounced in *USP37* KO cells, bringing the levels of RPA and γH2AX foci down to those observed in WT cells. Overall, these results suggested that USP37 is required for cells to overcome diverse forms of replication stress, and that in the context of topological stress caused by CPT or ICRF-193, its primary function is to counteract the action of TRAIP.

### USP37 prevents CMG disassembly by TRAIP upon TOP2α inhibition

Prior work showed that TRAIP ubiquitylates CMG^11,20–22^. We therefore explored whether USP37 counteracts TRAIP-dependent CMG ubiquitylation under conditions of topological stress. To this end, we used *Xenopus laevis* egg extracts, which recapitulate TRAIP-dependent CMG ubiquitylation when forks converge on an ICL^11^, as well as CRL2^Lrr1^-dependent CMG ubiquitylation when replisomes pass each other during termination^7^. We first confirmed that in this system, the TOP2 inhibitor ICRF-193 greatly delayed CMG unloading (Fig. 3a, compare lanes 1-3 and 7-9), as expected from the fact that it impairs replisome convergence (Extended Data Fig. 5a; Ref.^6^). In contrast with ICRF-193 treatment, TOP2α depletion (Extended Data Fig. 5b) had only a mild effect on CMG unloading (Extended Data Fig. 5c, lanes 1-3 vs. 7-9), commensurate with its more modest impairment of fork convergence that is probably still enabled by fork rotation or the action of accessory helicases^5,6,43,44^. Moreover, TOP2α immuno-depletion abrogated the strong effect of ICRF-193 on CMG unloading (Extended Data Fig. 5c, lanes 4-6 vs. 10-12). Together with the fact that TOP2α can accumulate ahead of the replisome (Extended Data Fig. 5d), these data strongly suggest that ICRF-193 delays fork convergence and CMG unloading primarily by trapping TOP2α complexes ahead of the replication fork.

**Fig. 3:**
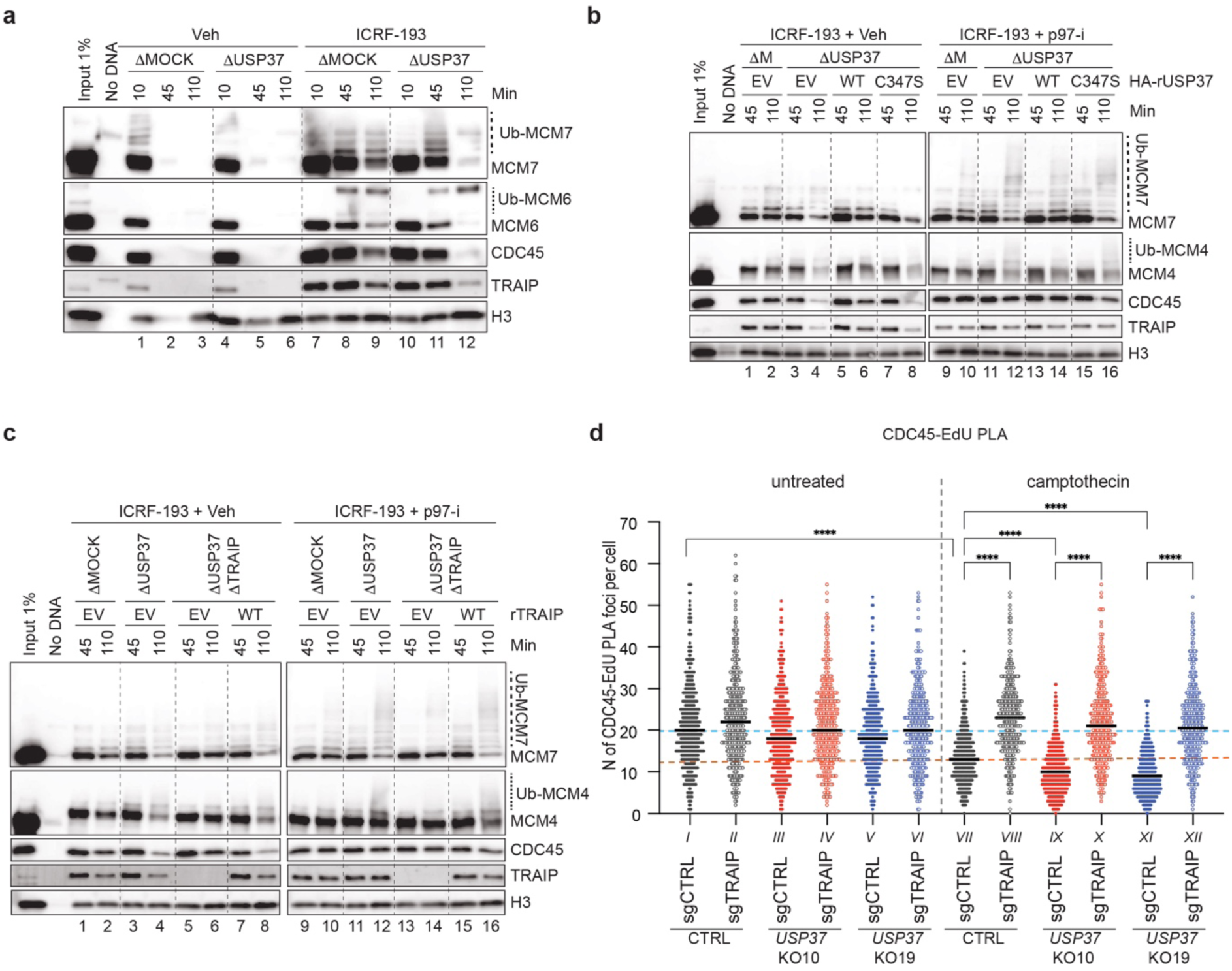
USP37 depletion induces premature CMG unloading by TRAIP in egg extracts treated with ICRF-193. **a,** Plasmid DNA was incubated in the indicated egg extracts in the presence or absence of 200 μM ICRF-193. At specified times, chromatin was recovered and immunoblotted for the indicated proteins. USP37 depletion efficiency is shown in Extended Data Fig. 5e. Ub-MCM7, ubiquitylated MCM7; Ub-MCM6, ubiquitylated MCM6. **b,** Plasmid DNA was incubated in the indicated egg extracts in the presence of 200 μM ICRF-193 and, where indicated, 200 μM NMS-873 (p97-i). Extracts were supplemented with recombinant WT or catalytically inactive (C347S) USP37 expressed in wheat germ extract, or wheat germ extract with empty vector (EV). At specified times, chromatin was recovered and immunoblotted for the indicated proteins. USP37 depletion efficiency and levels of recombinant proteins are shown in Extended Data Fig. 5f. ΔM, ΔMOCK; Ub-MCM4, ubiquitylated MCM4. **c,** As in **b**, but where indicated, TRAIP was co-depleted with USP37. Egg extracts were supplemented with recombinant WT TRAIP expressed in wheat germ extract, or wheat germ extract with EV. Depletion efficiencies and levels of recombinant proteins are shown in Extended Data Fig. 5g. **d,** Dot plot indicating the number of PLA foci between CDC45 and EdU (replicating DNA) in untreated or camptothecin treated cells. Bar represents median. n=3 independent experiments.

To test whether USP37 affects CMG unloading in these settings, we immuno-depleted USP37 from egg extracts (Extended Data Fig. 5e). While USP37 depletion did not influence CMG unloading during unperturbed replication (Fig. 3a, lanes 4-6), its absence accelerated CMG disassembly in the presence of ICRF-193 (Fig. 3a; compare lanes 7-9 and 10-12). Depending on extract preparation, CMG disassembly occurred between 90 and 120 min after replication initiation in USP37-depleted extracts. By contrast, USP37 depletion did not accelerate CMG unloading in extracts depleted of TOP2α, whether or not ICRF-193 was added (Extended Data Fig. 5c, lanes 7-12 vs. 13-18). Premature CMG unloading was suppressed by supplementing USP37-depleted extracts with wild-type HA-tagged USP37 (USP37^WT^; expressed in a transcription-translation extract), but not with catalytically inactive USP37 (USP37^C347S^; Fig. 3b, lanes 2, 4, 6, 8; Extended Data Fig. 5f). In the presence of the p97 inhibitor NMS-873 (p97-i), USP37 depletion led to polyubiquitylation of MCM7 and MCM6 on chromatin, which was most evident from loss of the unmodified forms of these proteins (Fig. 3b, lane 10 vs. 12). Furthermore, this effect was reversed by USP37^WT^ but not USP37^C347S^ (Fig. 3b, lanes 14 and 16). These results argue that when replisomes encounter trapped TOP2α complexes, USP37 suppresses CMG ubiquitylation and p97-dependent unloading.

We next tested whether CMG unloading in the absence of USP37 depends on TRAIP. Consistent with this idea, depletion of TRAIP (Extended Data Fig. 5g) inhibited premature CMG unloading in USP37-depleted extracts containing ICRF-193 (Fig. 3c, lanes 4 vs. 6). Moreover, compared to controls, TRAIP depletion increased the proportion of unmodified MCM7 (Fig. 3c, lane 12 vs. 14). Re-addition of TRAIP to such extracts rescued CMG unloading and ubiquitylation (Fig. 3c, lanes 8 and 16). We speculate that the remaining TRAIP-independent ubiquitylation of MCM7 (Fig. 3c, lane 14) is CRL2^Lrr1^-dependent and occurs due to termination of some forks despite the presence of ICRF-193 (Ref.^6^).

To address whether USP37 controls CMG unloading at sites of DNA replication in mammalian cells, we employed the SIRF (*in situ* Protein Interaction with Nascent DNA Replication Forks) assay^45^, which uses a fluorescent-based proximity ligation assay (PLA) to probe the proximity between a protein of interest and EdU-labelled replicating DNA. A PLA signal is only visible when an antibody against the protein of interest and an antibody against biotin, which detects biotinylated EdU, are located within 40 nm. When using the number of PLA foci as a readout for CDC45 localization at sites of DNA replication, we observed that CPT strongly reduced the number of PLA foci in control cells (conditions I vs. VII) and even further in *USP37* KO cells (compare VII vs. IX and XI) (Fig. 3d). Notably, TRAIP loss prevented CDC45 unloading in control and in *USP37* KO cells (conditions VIII, X, XII). Collectively, these results indicate that when forks experience stress in egg extracts and in mammalian cells, USP37 prevents CMG disassembly by TRAIP.

### USP37 suppresses TRAIP-dependent disassembly of converged CMGs

When TOP2α is inhibited in *Xenopus* egg extracts, converging replisomes stall several hundred base pairs (bp) apart, followed by slow fork merging^6^. We therefore hypothesized that in the presence of ICRF-193, TRAIP ubiquitylates CMGs that stall close to each other. To test this idea, we created a DNA substrate in which two tandem methylated HpaII DNA-protein cross-links (meDPCs) on the leading strands were placed 165 bp apart (Fig. 4a, top). Such tandem meDPCs cannot be degraded or bypassed by CMG and therefore pose a strong barrier to the replisome, preventing CMG unloading (Ref.^46^; Fig. 4a, lanes 1-2). However, when USP37 was depleted, the CMGs stalled at these tandem DPCs underwent unloading (Fig. 4a, compare lanes 2 and 4) and polyubiquitylation in the presence of p97-i (Fig. 4a, lanes 8 and 10). This CMG unloading and ubiquitylation was attenuated when TRAIP was also absent (Fig. 4a, lanes 6 and 12). These data thus indicated that USP37 suppresses replisome ubiquitylation and disassembly by TRAIP when CMGs stall at close range for an extended time.

**Fig. 4:**
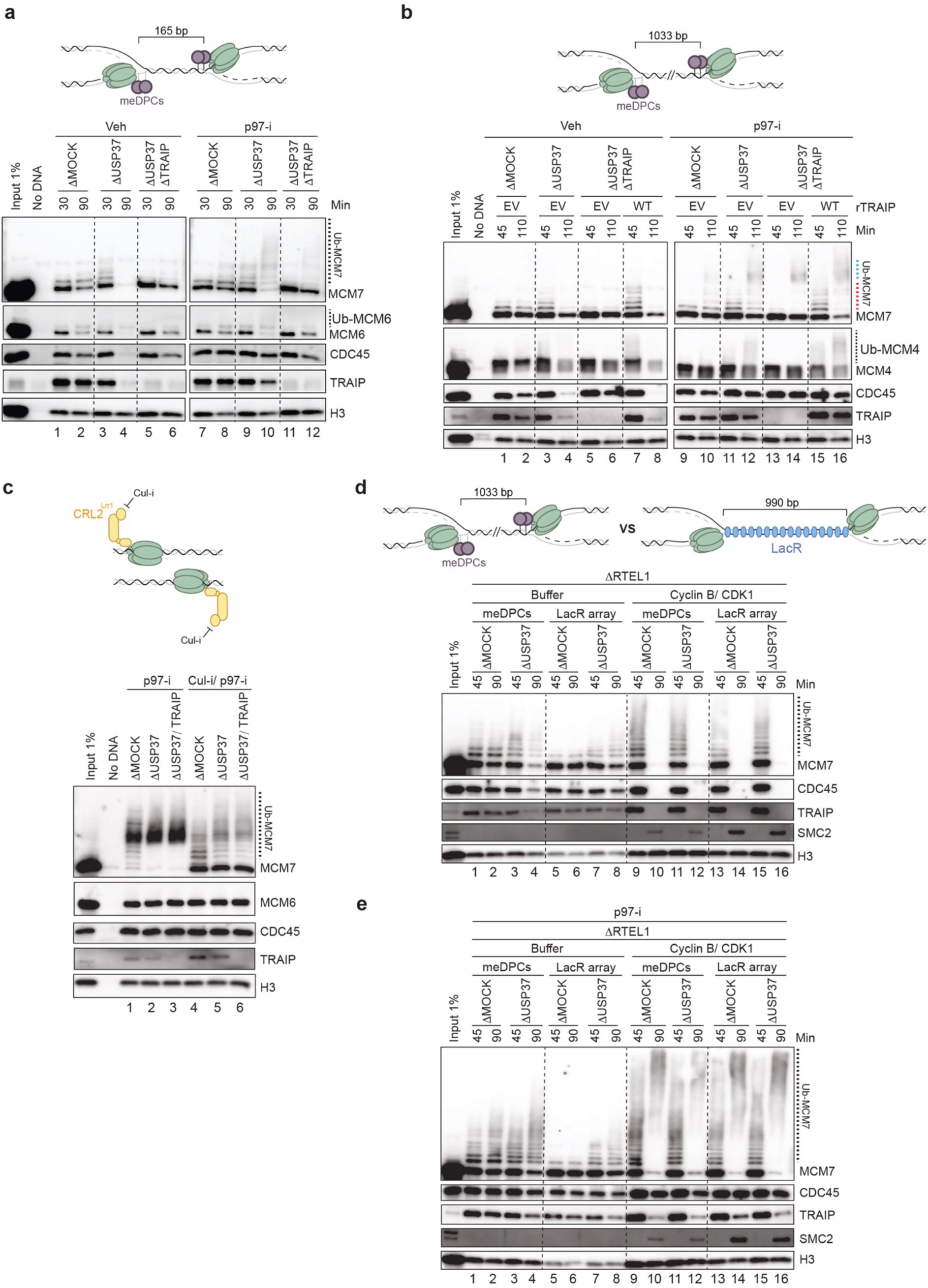
USP37 depletion induces TRAIP-dependent disassembly of CMGs converged at tandem DPCs, but not at terminated forks or ones separated by a LacR array. **a,** Top, a schematic of DNA protein cross-link substrate. meDPCs, methylated DNA protein cross-links. The square bracket indicates the distance between distal leading strand DPCs. Bottom, meDPCs substrate was incubated in indicated egg extracts in the presence or absence of 200 μM p97-i. At specified times, chromatin was recovered and immunoblotted for the indicated proteins. USP37 and TRAIP depletion efficiencies are shown in Extended Data Fig. 6a. **b,** Same as **a**, top, but the distance between distal leading strand DPCs was 1033 bp. Bottom, meDPCs substrate was incubated in the indicated egg extracts in the presence or absence of 200 μM p97-i. Extracts were supplemented with recombinant WT TRAIP expressed in wheat germ extract or wheat germ extract with empty vector (EV). Depletion efficiencies and levels of recombinant proteins are shown in Extended Data Fig. 6d. Dashed blue line indicates probable CRL2^Lrr1^-dependent ubiquitylation resulting from termination events. Dashed red line indicates TRAIP-dependent ubiquitylation. **c,** Top, a schematic showing terminated CMGs and inhibited CRL2^Lrr1^ by Cul-i. Bottom, plasmid DNA was replicated in the indicated egg extracts in the presence of p97-i (200 μM) and in the presence or absence of Cul-i (400 μM). At 90 min after replication initiation, chromatin was recovered and immunoblotted for the indicated proteins. USP37 and TRAIP depletion efficiencies are shown in Extended Data Fig. 6e. **d,** Top, a schematic of CMGs converged at the DNA protein cross-link substrate versus the LacR-bound *lacO* array. Bottom, plasmid DNA containing 32x *lacO* array was preincubated with LacR and then replicated in the indicated egg extracts. At 30 min after replication initiation, reactions were supplemented with either buffer or Cyclin B/CDK1 (50ng/ul). At the specified times, chromatin was recovered and immunoblotted for the indicated proteins. USP37 and RTEL1 depletion efficiencies are shown in Extended Data Fig. 6h. **e,** Same as **d**, but at 40 min after replication initiation, reactions were supplemented with p97-i (200 μM). USP37 and RTEL1 depletion efficiencies are shown in Extended Data Fig. 6i.

To explore the distance-dependence of CMG ubiquitylation, we generated a panel of meDPC substrates in which DPCs were separated by 56, 305, and 1033 bp (Extended Data Fig. 6b, top). All tested substrates led to stalling of converging CMGs in mock-depleted extracts, as seen from the persistence of CMG on chromatin at the late time point (Extended Data Fig. 6b, lanes 1-12). Strikingly, USP37 depletion enhanced CMG disassembly even when meDPCs were spaced ∼1000 bp apart (Extended Data Fig. 6b, lanes 1-12). Likewise, the ubiquitylation of MCM7 was similarly enhanced upon USP37 depletion in the context of all substrates (Extended Data Fig. 6b, lanes 13-24). Furthermore, in the context of USP37 depletion, the co-depletion of TRAIP blocked CMG ubiquitylation and unloading even when CMGs were 1 kb apart (Fig. 4b, lanes 3-4 vs. 5-6 and 11-12 vs. 13-14), and these effects were rescued by re-addition of TRAIP (Fig. 4b, lanes 8 and 16). Collectively, these results showed that in the absence of USP37, TRAIP can induce unloading of CMGs even when they are stalled at a considerable distance from each other.

### USP37 counteracts *trans* ubiquitylation of CMG by TRAIP

We showed previously that in S phase, TRAIP associates with the replisome and ubiquitylates proteins ahead of the replication fork “in *trans*” while being unable to ubiquitylate the CMG that it travels with “in *cis*” (Extended Data Fig. 1a; Ref.^11,12^). It was therefore unexpected that in USP37-depleted extracts, TRAIP was able to ubiquitylate CMGs that are stalled ∼1 kb apart. This observation raised the possibility that in the absence of USP37, TRAIP acquires the capacity to promote *cis* ubiquitylation of CMG (Extended Data Fig. 1b). To test this idea, we investigated whether, in the absence of USP37, TRAIP can ubiquitylate terminated CMGs that are normally only ubiquitylated by CRL2^Lrr1^ when CMGs have passed each other after fork convergence^11,21^. To this end, we replicated DNA in the presence of p97-i to prevent CMG unloading during termination, and we also added the general Cullin inhibitor MLN4924 (Cul-i) to minimize CRL2^Lrr1^-dependent ubiquitylation of CMG (Fig. 4c). As expected, Cul-i greatly reduced MCM7 ubiquitylation in mock-depleted extracts (Fig. 4c, lanes 1 and 4), but residual ubiquitylation was still observed, possibly due to incomplete inhibition of CRL2^Lrr1^ (Fig. 4c, lane 4) and/or the action of another ubiquitin E3 ligase. Although USP37 depletion enhanced this residual ubiquitylation of MCM7 (Fig. 4c, lane 5), most MCM7 remained unmodified, and importantly, the residual ubiquitylation was not dependent on TRAIP (Fig. 4c, lane 6). Moreover, in contrast to what was observed during CMG stalling during topological stress or at meDPCs, no ubiquitylation of MCM6 was observed in USP37-depleted extracts (Fig. 4c, lane 5). These data suggested that although TRAIP associates with terminated CMGs (Dewar et al. 2017), it is unable to support *cis* ubiquitylation, even in the absence of USP37. This result further implied that in the absence of USP37, TRAIP promotes ubiquitylation of distant CMGs in *trans*.

We hypothesized that co-localization of stalled CMGs may be required for TRAIP-mediated *trans* ubiquitylation in the absence of USP37. To test this idea, we employed a DNA substrate containing an array of 32 *lac* operator (*lacO*) repeats (comprising 990 bp) to which we bound the *lac* repressor (LacR). LacR-arrays slow CMG progression and delay its unloading (Ref.^21,47^, Fig. 4d, lanes 5-6), and they are expected to disrupt co-localization of replisomes by stalling CMGs at the outer edges of the *lacO* array (Fig. 4d, top). To minimize CMG unloading events due to slow replisome progression through the LacR barrier, we depleted the RTEL1, an accessory helicase that normally helps CMG pass through proteinaceous barriers (Extended Data Fig. 6f and Ref.^12,46^). When the LacR-bound plasmid DNA was replicated in egg extract immuno-depleted of both USP37 and RTEL1, stalled CMGs were not unloaded (Fig. 4d, lanes 5-8), and they underwent only a very low level of ubiquitylation (Fig. 4e, lanes 5-8). This contrasted with results obtained with the DPC-containing substrate in the same extract (Fig. 4d, lanes 1-4 and Fig. 4e, lanes 1-4). However, in mitotic extracts, CMGs stalled at DPCs or *lacO* array were unloaded and hyper-ubiquitylated with similar efficiencies regardless of USP37 absence (Fig. 4d, lanes 9-16 and Fig. 4e, lanes 9-16) suggesting that hyperactivation of TRAIP in mitosis occurs via a different mechanism and doesn’t require co-localization of replisomes. Collectively, these results are consistent with a model in which TRAIP-mediated CMG ubiquitylation at long-range in interphase involves a *trans* mechanism that is mediated by co-localization of replisomes.

### Evidence that USP37 functions by binding to CDC45

Previous experiments suggested that USP37 may be a component of the replisome^7,36,37^. However, we did not detect USP37 on replicating DNA in plasmid pull-down experiments in egg extracts (Extended Data Fig. 7a). We speculated this is because plasmid pull-down is a relatively lengthy procedure during which loosely bound proteins dissociate from chromatin^48^. To circumvent this potential issue, we used a rapid sperm chromatin isolation assay^49^. In this setting, we detected USP37 on chromatin, as seen previously^7^, and this binding was abolished by the replication-licensing inhibitor geminin and by inhibitors of CDK and DDK kinases, which are required to convert pre-replication complexes into active CMG helicases (Fig. 5a). We thus concluded that USP37 loads onto chromatin in a manner dependent on CMG assembly.

**Fig. 5:**
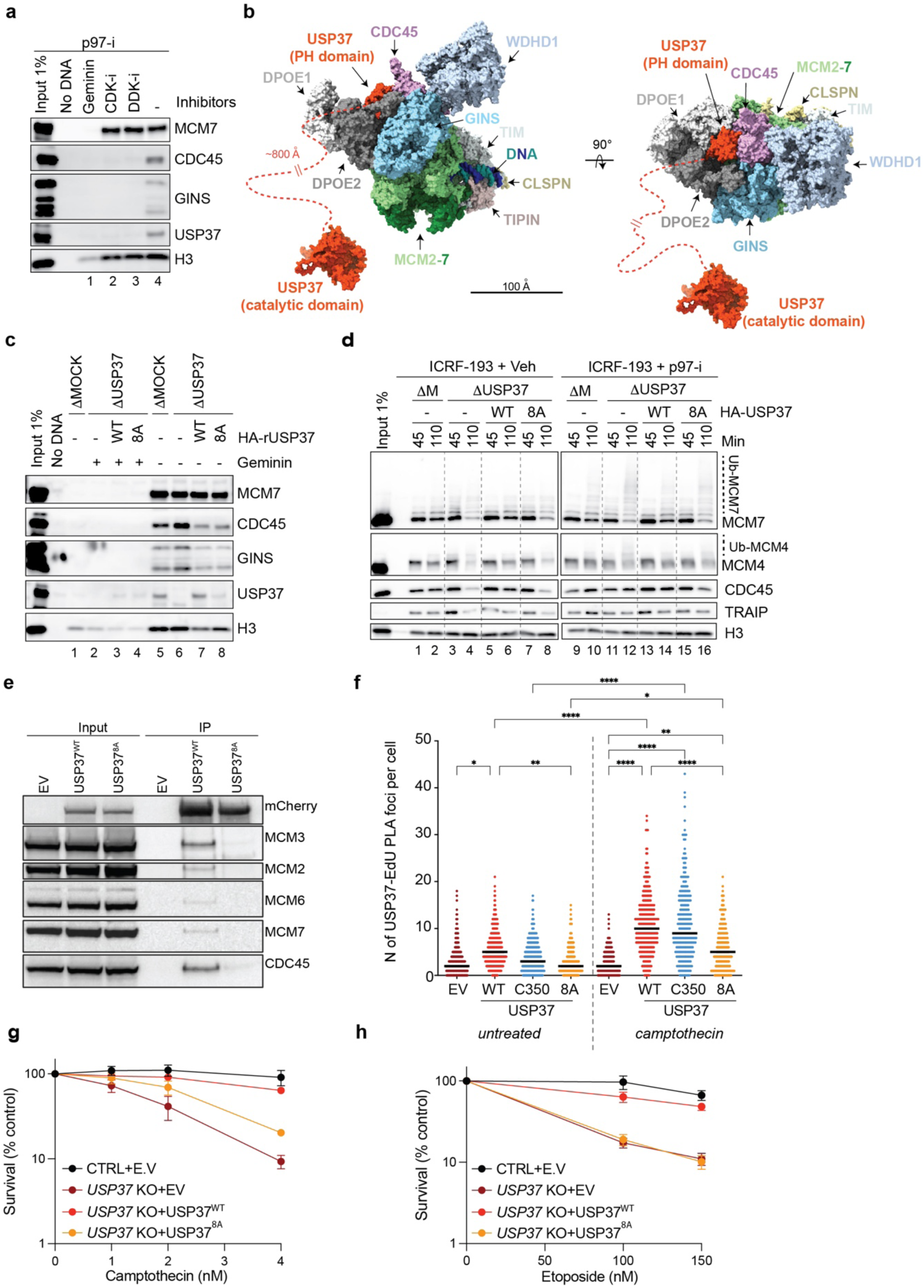
Predicted CDC45-USP37 interaction surface is important for USP37 function at the replisome. **a,** *Xenopus* sperm chromatin was incubated in egg extracts supplemented with indicated inhibitors for 10 min, then recovered and blotted for the indicated proteins. p97-i, NMS-873; CDK-i, p27Kip; DDK-i, PHA-767491. **b,** Right, the predicted structure in Extended Data Fig. 7c was aligned to the Cryo-EM structure of human CMG (PDB: 7pfo). Left, the same model rotated by 90°. Orange dashed line indicates a flexible region connecting the PH domain of USP37 and the catalytic domain. **c,** *Xenopus* sperm chromatin was incubated in mock- or USP37-depleted egg extracts supplemented with recombinant WT or putative CDC45-binding USP37 mutant (8A) expressed in wheat germ extract or wheat germ extract containing empty vector (-), then recovered and blotted for the indicated proteins. DNA-replication inhibitor, Geminin was added where indicated to monitor non-specific binding. See also Extended Data Fig. 7k for depletion efficiencies and recombinant protein levels. **d,** Plasmid DNA was incubated in the indicated egg extracts in the presence of 200 μM ICRF-193 and, where indicated, 200 μM NMS-873 (p97-i). Extracts were supplemented with the indicated recombinant USP37 variants expressed in wheat germ extract or wheat germ extract containing empty vector (EV). At specified times, chromatin was recovered and blotted for the indicated proteins. USP37 depletion efficiencies and levels of recombinant proteins are shown in Extended Data Fig. 7l. **e,** Extracts of HEK293T cells expressing mCherry (empty vector, EV) or mCherry-USP37^WT^ or mCherry-USP37^8A^ were subjected to mCherry immunoprecipitation (IP) followed by western blotting for the indicated proteins. This experiment was repeated three times with similar results. **f,** Dot plot indicating the number of PLA foci between mCherry-tagged E.V, USP37^WT^, USP37^C3^^50^, or USP37^8A^ and EdU (replicating DNA) in untreated or camptothecin treated cells. Bar represents median. n=3 independent experiments. **g-h,** Clonogenic survival assays of control (CTRL) cells or USP37 KO (clone 10) cells complemented with vectors expressing mCherry (EV), mCherry-USP37^WT^ or mCherry-USP37^8A^ (defective for CDC45 interaction) upon treatment with camptothecin (**g**) or etoposide (**h**). Note that CTRL+EV and USP37 KO+ mCherry-USP37WT samples are the same as in Fig. 1h and i. n=3 independent experiments. Bars represent means ± SEM.

We next explored how USP37 associates with replisomes by using AlphaFold-Multimer (AF-M) to screen *in silico* for potential USP37 binding partners among 285 proteins involved in genome maintenance (predictomes.org; Ref.^50^). We scored the results using a classifier called SPOC (structure prediction and omics classifier) that is trained to distinguish between high and low confidence AF-M predictions^51^. According to this metric, cohesin subunits (STAG2, STAG1, and SMC3) as well as replication factors, RPA2 and CDC45, were the top-ranked proteins predicted to bind USP37 (Supplementary Table 1). Since USP37 interaction with cohesin was reported previously^37^, we tested whether immunodepletion of SMC3 phenocopies USP37 depletion in causing premature CMG unloading. However, depletion of SMC3 (Extended Data Fig. 7b) had no measurable effect on CMG unloading at meDPCs (Extended Data Fig. 7c, compare lanes 2,4, and 6), arguing against any role for cohesin in recruiting USP37 to the replisome. Since USP37 still associates with terminated replisomes in the absence of ssDNA^7^, RPA2 was also unlikely to recruit USP37. We therefore focused on the core replisome component, CDC45, which ranked fourth among potential interactors. Conversely, USP37 was ranked second among 285 genome maintenance proteins predicted to bind CDC45, ahead of GINS1 (Supplementary Table 1 and Extended Data Fig. 7d, e). Similar results were obtained when *Xenopus* USP37 was folded *in silico* with *Xenopus* replication proteins (Supplementary Table 1). AF-M predicted that an antiparallel β-sheet of the USP37 N-terminal pleckstrin homology (PH) domain binds CDC45 adjacent to DNA polymerase epsilon subunits DPOE1 and DPOE2 (Extended Data Fig. 7f-h) without clashing with any known replisome proteins (Fig. 5b). Furthermore, consistent with the USP37-CDC45 interaction having an important function, the predicted binding surface of USP37, as well as the interacting residues, are highly conserved between human and *Xenopus* (Extended Data Fig. 7g-i). Because the USP37 catalytic domain is connected to the PH domain via an approximately 800 Å unstructured region, USP37 should be able to de-ubiquitylate most components of the replisome (Fig. 5b).

To test the function of the predicted USP37-CDC45 interaction, we mutated eight highly conserved USP37 residues at the predicted interface to alanine, generating USP37^8A^ (Extended Data Fig. 7i). The catalytic activity of USP37^8A^ was minimally affected by these changes, as monitored by its ability to cleave K48-linked tetraubiquitin at similar levels as the unmutated control protein (Extended Data Fig. 7j). By contrast with the wild-type protein, USP37^8A^ did not bind chromatin efficiently (Fig. 5c, lane 8), and it failed to prevent premature CMG ubiquitylation and unloading in the presence of ICRF-193 in egg extract (Fig. 5d, lane 6 vs. 8 and 14 vs. 16). Furthermore, in mammalian cells, USP37^WT^ but not USP37^8A^ co-precipitated with CDC45 and other CMG components (Fig. 5e). To further confirm USP37 localization to the replisome in mammalian cells, we employed the SIRF assay. By using the number of PLA foci as an indicator of USP37 localization at replicating DNA, we detected a basal level of USP37 at sites of DNA replication in untreated conditions. Upon treatment with CPT, wild-type (USP37^WT^) and catalytically inactive (USP37^C3^^50^) variant localization at replicating DNA increased, while USP37^8A^ did so to a much lesser extent (Fig. 5f). Finally, to monitor whether the USP37-CDC45 interaction is important, we monitored the sensitivity to genotoxic agents of *USP37* KO cells complemented with an empty vector (E.V), USP37^WT^, or USP37^8A^. USP37^WT^ but not USP37^8A^ complemented the CPT and etoposide hypersensitivities of *USP37* knockout cells (Fig. 5g-h, Extended Data Fig. 2i, and Extended Data Fig. 8a-b). Collectively, these data suggested that USP37’s interaction with CDC45 counteracts TRAIP-mediated CMG ubiquitylation, which mitigates the toxic effects of topoisomerase inhibitors.

## Discussion

Replisomes are disassembled during replication termination and at ICLs. However, before termination, and at most types of replication obstacles, replisome disassembly is deleterious. Yet, how cells prevent inadvertent loss of the replisome is incompletely understood. Based on experiments in cell-free extracts and mammalian cells, we report here that under certain forms of replication stress, USP37 prevents CMG ubiquitylation and unloading by the TRAIP E3 ubiquitin ligase. We propose that this regulation is critical to allow the completion of DNA replication and the suppression of genome instability.

We uncovered a complex genetic interaction between *USP37* and *TRAIP*. While *USP37* knockout cells were hypersensitive to TOP1 and TOP2 inhibitors, this phenotype was ameliorated in *TRAIP*/*USP37* double knockouts, indicative of genetic suppression or “synthetic rescue”. Consistent with the cell viability data, we observed that the increased number of CPT-induced RPA and γH2AX foci in *USP37* knockout cells was mitigated by TRAIP loss, indicating that USP37 protects against the accumulation of ssDNA and DSBs resulting from aberrant TRAIP activity. The simplest interpretation of these results is that in the absence of USP37, hyper-ubiquitylation of one or more TRAIP substrates leads to DNA damage and cell death. Conversely, the CPT sensitivity in *TRAIP* knockouts was also largely suppressed by ablating *USP37*, demonstrating synthetic rescue in the other direction. Similarly, while our genetic screens revealed that TRAIP loss reduced fitness of WT cells in the absence of external stress, as observed previously^13,40^, in USP37-deficient cells, the loss of TRAIP had no further deleterious effect. These results suggest that TRAIP has important targets whose ubiquitylation is critical for cell viability in the face of endogenous and exogenous stress. We speculate that when USP37 is lost in the context of TRAIP deficiency, a different E3 ubiquitin ligase whose activity is normally restrained by USP37, can take over TRAIP’s function. Thus, the interplay of TRAIP and USP37 is complex, and we focus here on the toxic activity of TRAIP in the absence of USP37. Notably, TRAIP mutations are associated with microcephalic dwarfism, a condition characterized by diminished growth and reduced brain size^52^. Thus, we speculate that inactivating USP37 or inhibiting its activity might ameliorate phenotypes associated with this condition.

Our observations in *Xenopus* egg extracts provide an attractive explanation for the synthetic rescue of hypersensitivity to topoisomerase inhibitors observed in *USP37/TRAIP* double-knockout cells compared to *USP37* knockout cells. In this cell-free system, resolution of topological stress is essential to allow rapid replisome convergence and unloading^6^. Notably, we have found that when replisome convergence was delayed by the TOP2 inhibitor ICRF-193, USP37 was required to suppress TRAIP-dependent premature CMG unloading. Similarly, when we stalled forks with DNA-protein cross-links, USP37 prevented TRAIP-dependent CMG unloading. Based on these observations, we propose that when converging forks stall, USP37 prevents CMG unloading by TRAIP (Fig. 6), and we speculate that a similar phenomenon occurs when mammalian cells are treated with topoisomerase inhibitors.

**Fig. 6:**
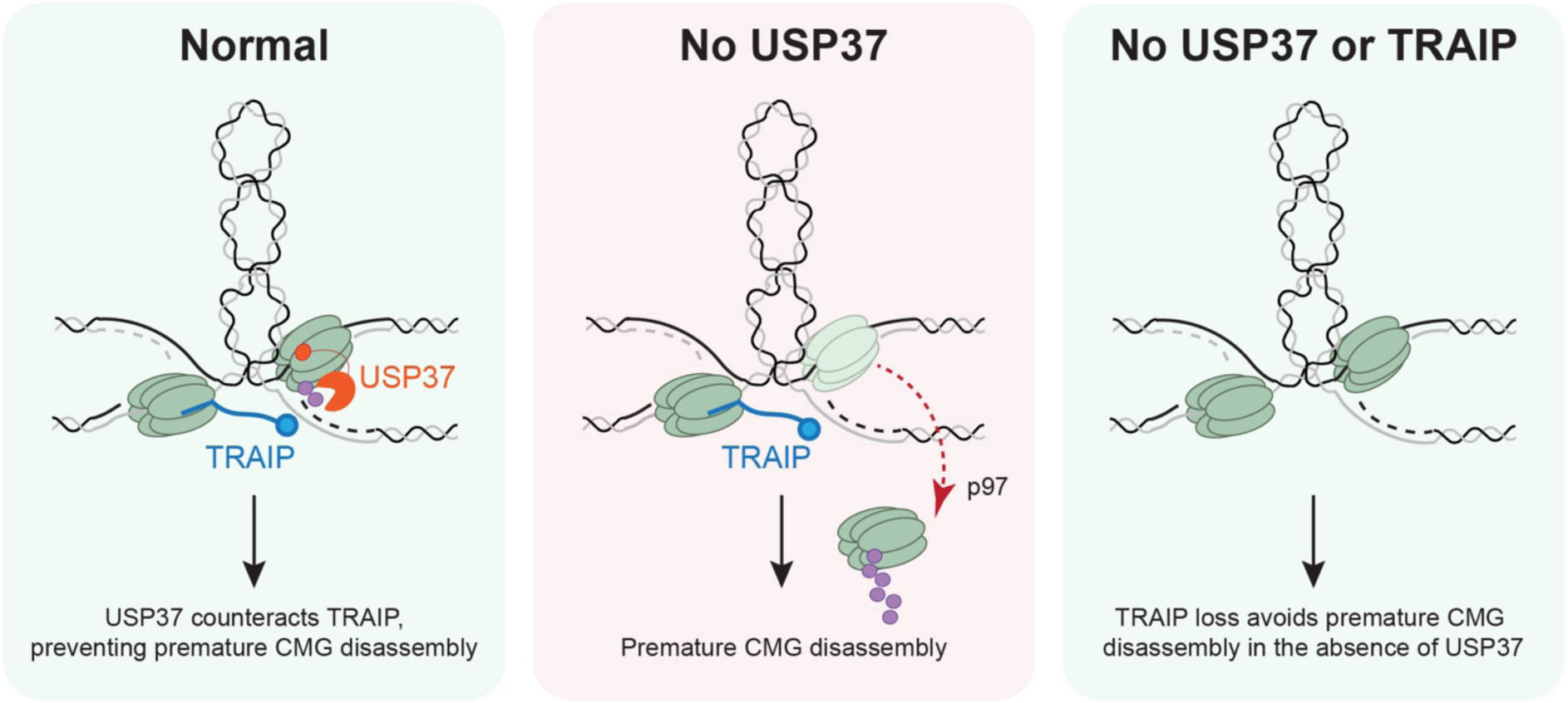
Model of USP37’s role in protecting replisomes from premature disassembly In wild-type cells (WT), USP37 counteracts TRAIP ubiquitylation when CMGs are stalled at sites of DPCs/topological stress. Thus, USP37 activity allows replisomes to eventually complete DNA synthesis and be unloaded by a termination-specific CRL2^Lrr1^-dependent pathway. In the absence of USP37, TRAIP hyper-ubiquitylates CMG at sites of topological stress, causing premature CMG disassembly, DNA damage, and cell death. When both USP37 and TRAIP are absent converging CMGs cannot be prematurely ubiquitylated and, thus, may ultimately terminate normally. For simplicity, the region in between two stalled replisomes is shown as plectonemes.

We also found that TRAIP promotes CMG unloading in USP37-depleted extracts even when forks were stalled ∼1 kb apart. This observation is surprising because we previously showed that in interphase, TRAIP normally only ubiquitylates proteins ahead of the fork (*trans* ubiquitylation) while being unable to ubiquitylate the CMG with which it travels (*cis* ubiquitylation). How can TRAIP ubiquitylate CMGs separated by 1 kb in the absence of USP37? While *cis* ubiquitylation occurs in mitosis, we do not believe that USP37 suppresses *cis* ubiquitylation in interphase because even in USP37-depleted extracts, TRAIP was not able to ubiquitylate terminated replisomes that passed each other. Instead, we hypothesize that when forks stall at DPCs positioned at a distance from one another, positive supercoiling, chromatin compaction, and/or physical coupling of replisomes bring the CMGs close enough together to allow *trans* ubiquitylation by TRAIP. In fact, the spatial and functional interaction of two converging replisomes has been proposed to occur constantly in S phase in human and mouse cells^53^. Consistent with this interpretation, we found that USP37 did not induce de-ubiquitylation of CMGs stalled on the outer edges of a 1 kb LacR array, which would probably disrupt 3D chromatin architecture and/or physical coupling of replisomes. Although LacR is a non-covalent protein barrier that allows slow CMG progression (Extended Data Fig. 6f), in the absence of RTEL1, we observed that fork progression through the array was minimal. Therefore, we conclude that LacR and tandem DPCs should have a similar effect on replisome progression, implying that the difference in CMG ubiquitylation in these two settings is related to disruption of CMG co-localization.

Unlike CMGs stalled at long range, replicative helicases stalled at ICLs undergo extensive ubiquitylation by TRAIP even in the presence of USP37^11^. Thus, an important question is how TRAIP overcomes the inhibitory activity of USP37 in this setting? We envision two explanations: first, TRAIP activity is dramatically higher at close range, such that it overwhelms USP37 activity at converged CMGs; second, USP37 is negatively regulated when forks meet on either side of an ICL. More work will be required to unravel how the balance of TRAIP and USP37 is regulated under different conditions.

Although aphidicolin induces DNA damage and cell death in *USP37* knockout cells, TRAIP loss did not rescue these phenotypes, suggesting that USP37 counteracts the activity of another E3 ubiquitin ligase in this setting. In line with this observation, we found that TRAIP loss on its own did not hypersensitize cells to aphidicolin, and aphidicolin induced complete TRAIP dissociation from chromatin in egg extracts (Extended Data Fig. 5d). As such, TRAIP is not available to ubiquitylate proteins at the replication fork during this form of stress. As the only other factor known to ubiquitylate CMG, CRL2^Lrr1^ is a strong candidate for the ligase counteracted by USP37 under such circumstances.

Together with previous studies^7,36,37^, our findings strongly suggest that USP37 is a constitutive replisome component. Consistent with this idea, we have established that USP37 binding to chromatin requires DNA licensing and the activities of the DDK and CDK kinases, demonstrating that USP37 recruitment requires CMG assembly. Moreover, AlphaFold made strong reciprocal predictions that USP37 and CDC45 interact, highlighting CDC45 as a likely tethering point for USP37 at the replisome. Accordingly, we found that mutations in the USP37 PH domain predicted to disrupt CDC45 binding impaired association of USP37 with replicating chromatin and its ability to deubiquitylate stalled CMGs in egg extracts. Additionally, this USP37 mutant failed to co-immunoprecipitate with CMG, and it failed to protect cells from TOP1 and TOP2 poisons. We speculate that the presence of a long flexible linker between the PH and catalytic domains of USP37 allows this DUB to deubiquitylate multiple proteins at the fork (Fig. 5b). Consistent with this, we observed enhanced ubiquitylation of multiple CMG subunits upon USP37 loss (MCM7, MCM4 and MCM6). Whether USP37 can also deubiquitylate and stabilize proteins ahead of the replication fork is an interesting subject for future investigation.

In conclusion, our findings identify USP37 as a regulator of genome integrity that prevents premature disassembly of stressed replisomes by TRAIP. Furthermore, in light of the observation that the catalytic activity of USP37 is essential to protect against topoisomerase inhibitors and replication stalling agents, our work suggests that USP37 inhibitors might be selectively toxic to cancer cells exhibiting elevated replication stress. Furthermore, our results suggest that inactivating USP37 or inhibiting its activity might ameliorate phenotypes associated with microcephalic dwarfism, which is caused by mutations in TRAIP^52^.

## Acknowledgements

We thank James Dewar, Karim Labib, and members of the Walter and Jackson laboratories for critical feedback on the manuscript. We also acknowledge all members of the Cancer Research UK Cambridge Institute facilities for assistance. J.C.W. is supported by NIH grant HL098316. J.C.W. is a Howard Hughes Medical Institute Investigator and an American Cancer Society Research Professor. Research in the S.P.J laboratory is supported by Cancer Research UK (CRUK) Discovery Award DRCPGM\100005, CRUK core grant A:29580 and ERC Synergy Award 855741 (DDREAMM). S.P.J is an employee of the University of Cambridge. GD’A and A.V. are funded by an ERC Synergy Award 855741. This project has received funding from AIRC and the European Union’s Horizon 2020 research and innovation programme under Marie Skodowska-Curie grant agreement no. 800924 to G.D.A. Research in the G.S.S. laboratory is supported by a CRUK programme grant (C17183/A23303).

## Contributions

O.V.K. performed all experiments in *Xenopus* egg extracts and also generated Fig. 5b and Extended Data Fig. 7d – i. G.D’A. performed experiments in Fig. 1, 2, 3e, 3g-h, Extended Data Fig. 2-4, 8, with the help of A.V., D.P., and S.L.R. AF-M predictions were performed by E.S. and O.V.K. Large-scale prediction screening in Supplementary Table 1 and analysis were performed by E.S. Fig. 3d and 5f were generated by M.R.G. and S.S.J. under the supervision of G.S.S. R.A.W. generated pAW18 vector and validated recombinant TRAIP activity in egg extracts. O.V.K., G.D’A., J.C.W., and S.P.J. wrote the manuscript.

---

**Extended Data Fig. 1:**
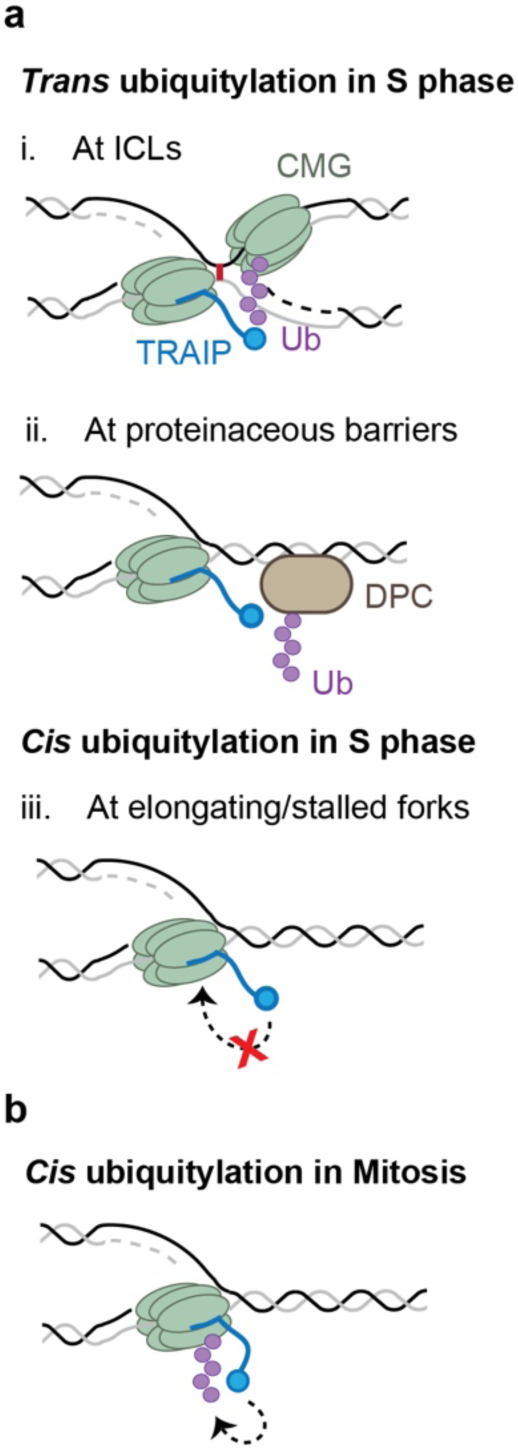
Model for TRAIP function in S and M phases. **a,** Using its *trans*-ubiquitylation mode in S phase, TRAIP can ubiquitylate CMGs that have converged on an ICL (i) or DPCs and other obstacles encountered by the replisome (ii), but not the replisome it travels with (which we call *cis* ubiquitylation) (iii). **b,** In mitosis, TRAIP undergoes a conformational change that allows it to *cis* ubiquitylate the CMG it travels with.

**Extended Data Fig. 2:**
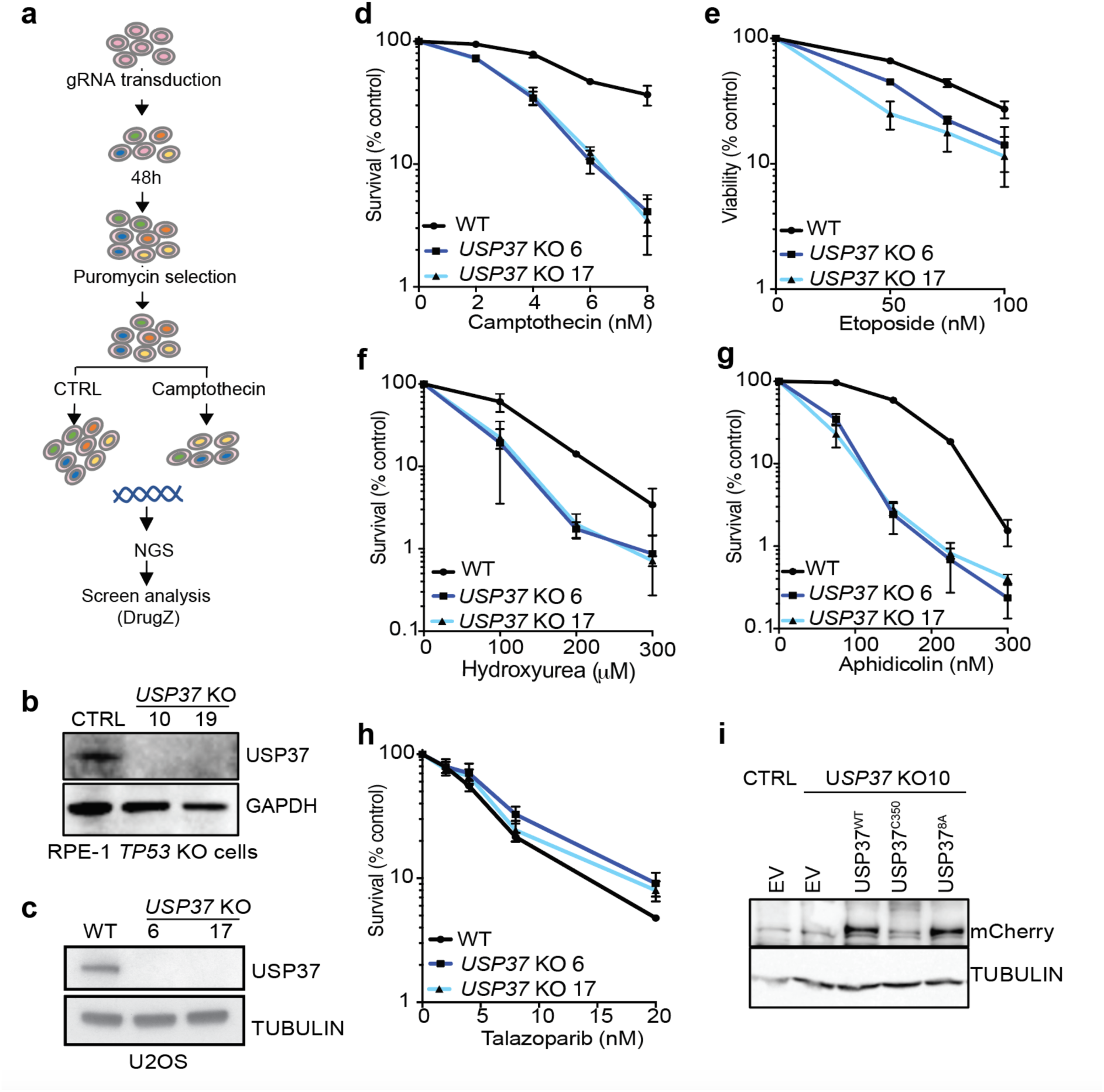
*USP37* knock-out hypersensitises to topoisomerase inhibitors and replication stress inducing agents. **a,** Schematic of the CRISPR screen in U2OS cells. **b-c,** Western blot validation of *USP37* KO in RPE-1 *TP53* KO cells (**b**) and U2OS cells (**c**). **d-h,** Clonogenic survival assays of WT and *USP37* KO U2OS cells upon treatment with (**d**) camptothecin, (**e**) etoposide, (**f**) hydroxyurea, (**g**) aphidicolin or (**h**) talazoparib. n ≥2 independent experiments. Bars represent means ± SEM. **i,** Western blot validation of mCherry-USP37^WT^, mCherry-USP37^C350A^ (catalytically inactive), or mCherry-USP37^8A^ expression in *USP37* knockout cells.

**Extended Data Fig. 3:**
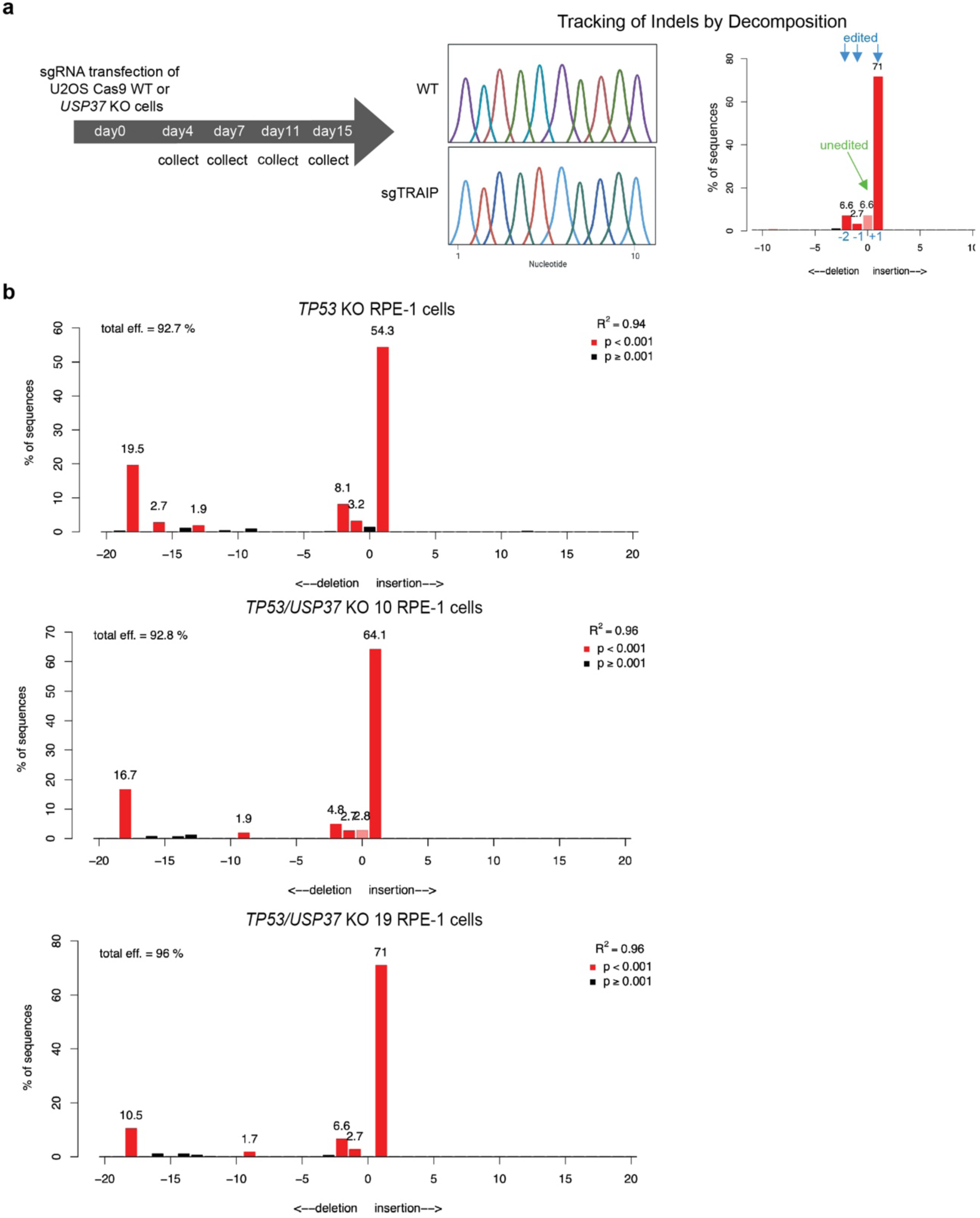
Validation of *TRAIP* KO in *TP53* KO and *TP53/USP37* KO RPE-1 cells. **a,** Scheme of the tracking indels by decomposition (TIDE)-based cell competition assay shown in Fig. 2b. Scheme generated with BioRender. **b,** TIDE-validation of *TRAIP* KO in *TP53* KO and *TP53/USP37* KO RPE-1 cells.

**Extended Data Fig. 4:**
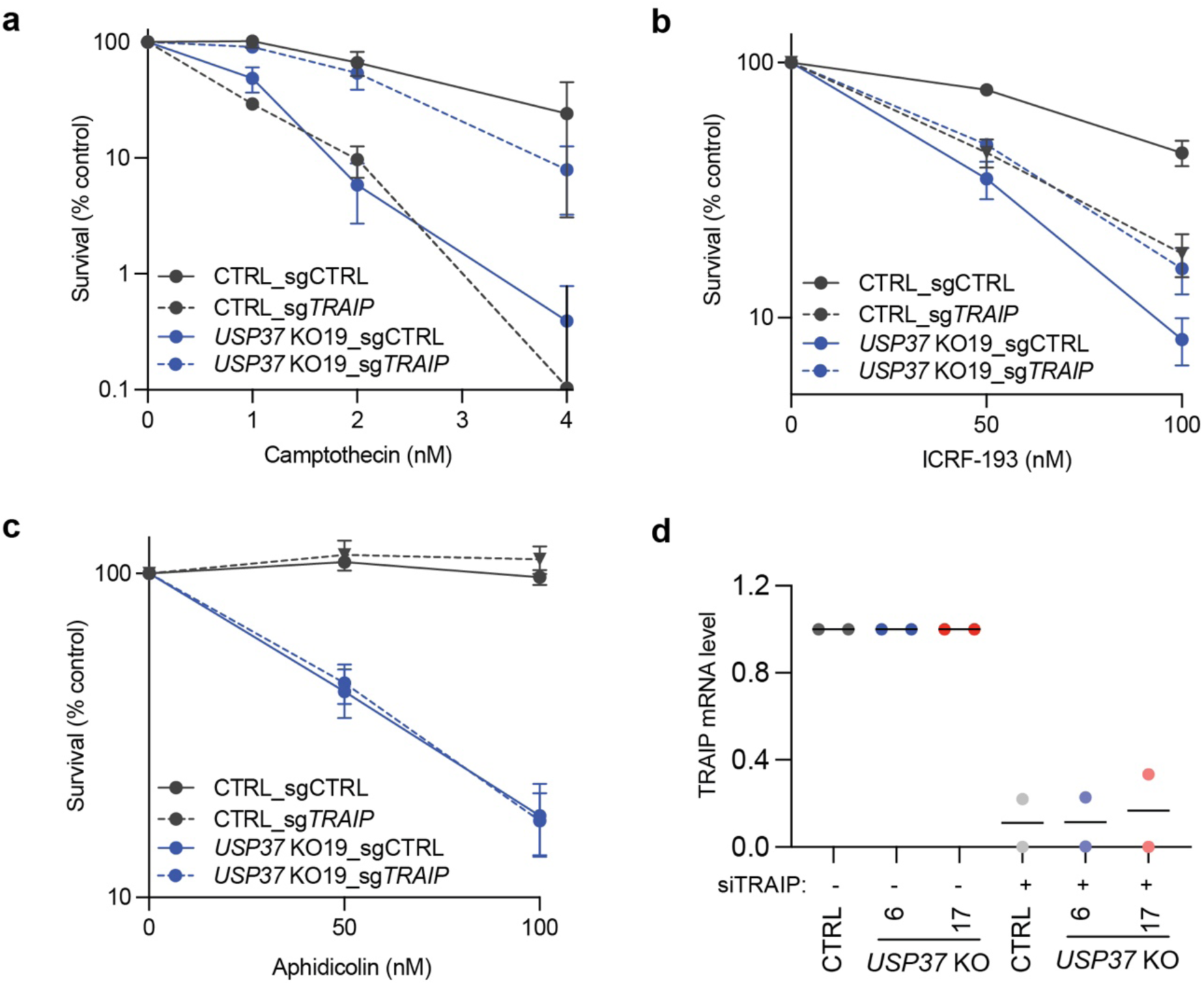
TRAIP loss improves the viability of *USP37* KO cells upon treatment with topoisomerase inhibitors but not aphidicolin. **a-c,** Clonogenic survival assays as in Fig. 2c-e of CTRL or a different *USP37* KO clone transduced with a control sgRNA (LacZ) or with a sgRNA targeting *TRAIP* upon treatment with camptothecin (**a**), ICRF-193 (**b**) or aphidicolin (**c**). The CTRL data are the same as in Fig. 2c-e. n=3 independent experiments. Bars represent means ± SEM. Half plot points in **c** indicate zero percent viability. **d,** RT-qPCR experiment to monitor TRAIP knock-down efficiency. n=2 independent experiments. Bars represent means.

**Extended Data Fig. 5:**
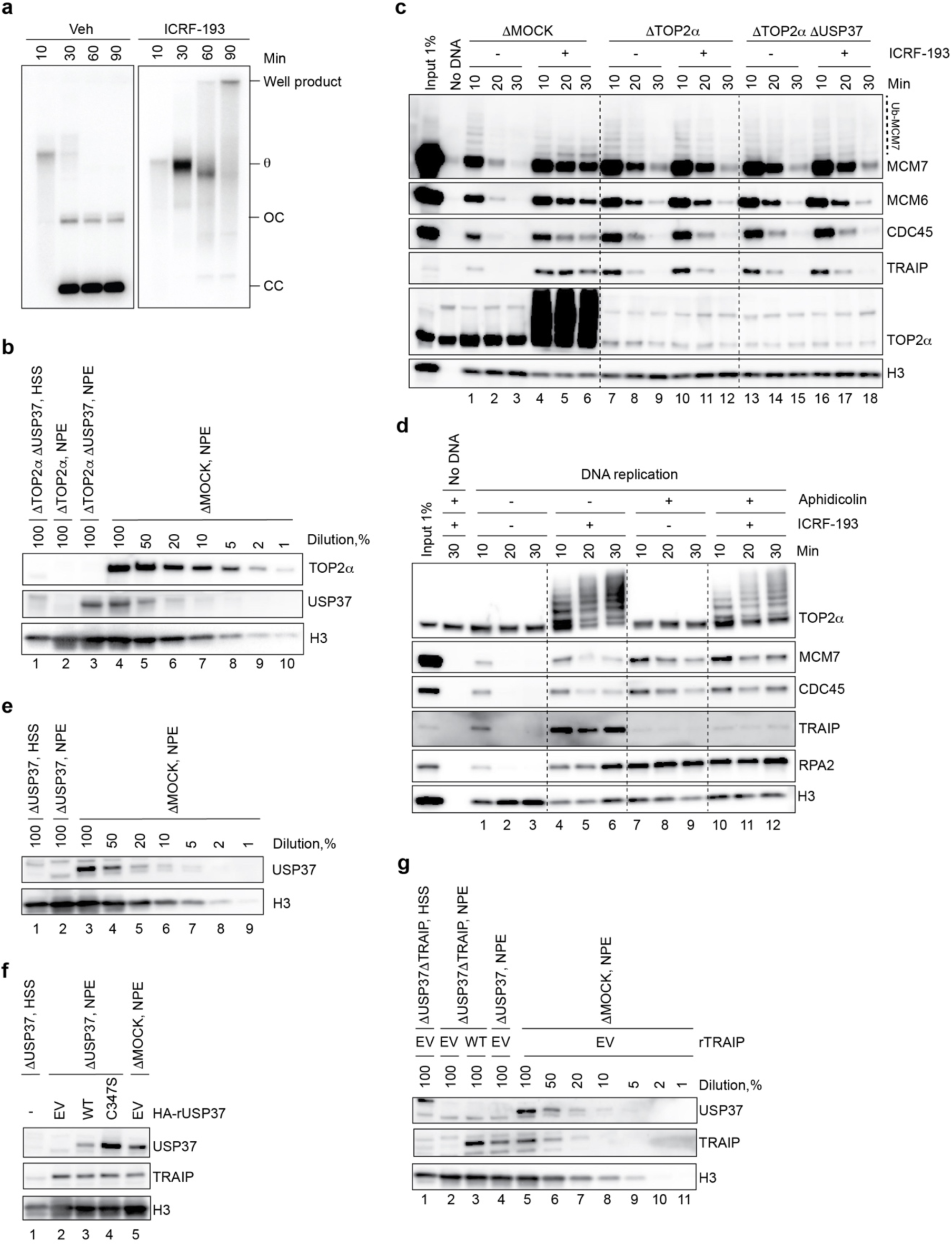
CMG ubiquitylation by TRAIP during topological stress requires trapping of TOP2α on DNA. **a,** Plasmid DNA was replicated in the presence or absence of 200 μM ICRF-193 in extracts containing [α-^32^P]dATP. Replication intermediates were then separated on a native agarose gel and visualized by autoradiography. At early time points, replication of plasmid DNA generates a “theta” structure (θ), a late replication intermediate formed when replisomes converge, that is subsequently converted to fully replicated closed and open circular products (“CC” and “OC”, respectively). ICRF-193 delays conversion of theta structures to CC and OC, indicative of impaired replisome convergence, and causes accumulation of a smear that probably represents highly catenated daughter molecules (Ref.^6^, see Extended Data Fig. 2a). The images are separate agarose gels, which were processed and imaged in parallel. **b,** Western blot analysis of mock, USP37, and TOP2α depletions. Related to Extended data Fig. 5c. **c,** Plasmid DNA was incubated in the indicated egg extracts in the presence or absence of 200 μM ICRF-193. At the specified times, chromatin was recovered and blotted for the indicated proteins. **d,** Plasmid DNA was incubated in the indicated egg extracts in the presence or absence of 200 μM ICRF-193 and 50 ng/μl aphidicolin. At specified times, chromatin was recovered and immunoblotted for the indicated proteins. Aphidicolin inhibits DNA synthesis and thus prevents formation of pre-catenanes while still allowing DNA unwinding and accumulation of supercoils^8,54,55^, which are presumably located ahead of the fork. As expected, aphidicolin increased RPA binding and decreased TOP2α recruitment to chromatin (lane 7 vs 1)^6^. However, in the presence of aphidicolin, ICRF-193 still caused accumulation of TOP2α on DNA, albeit to a lesser extent (lanes 10-12 vs 4-6) suggesting that TOP2α can be trapped on supercoils ahead of the replisome. **e,** Western blot analysis of mock and USP37 depletions. **f,** Western blot analysis of mock and USP37 depletions supplemented with recombinant USP37 expressed in wheat germ extract. Abbreviations as in Fig. 3b. **g,** Western blot analysis of mock, USP37, and TRAIP depletions supplemented with recombinant TRAIP expressed in wheat germ extract. Abbreviations as in Fig. 3c. Blue arrows, endogenous TRAIP.

**Extended data Fig. 6:**
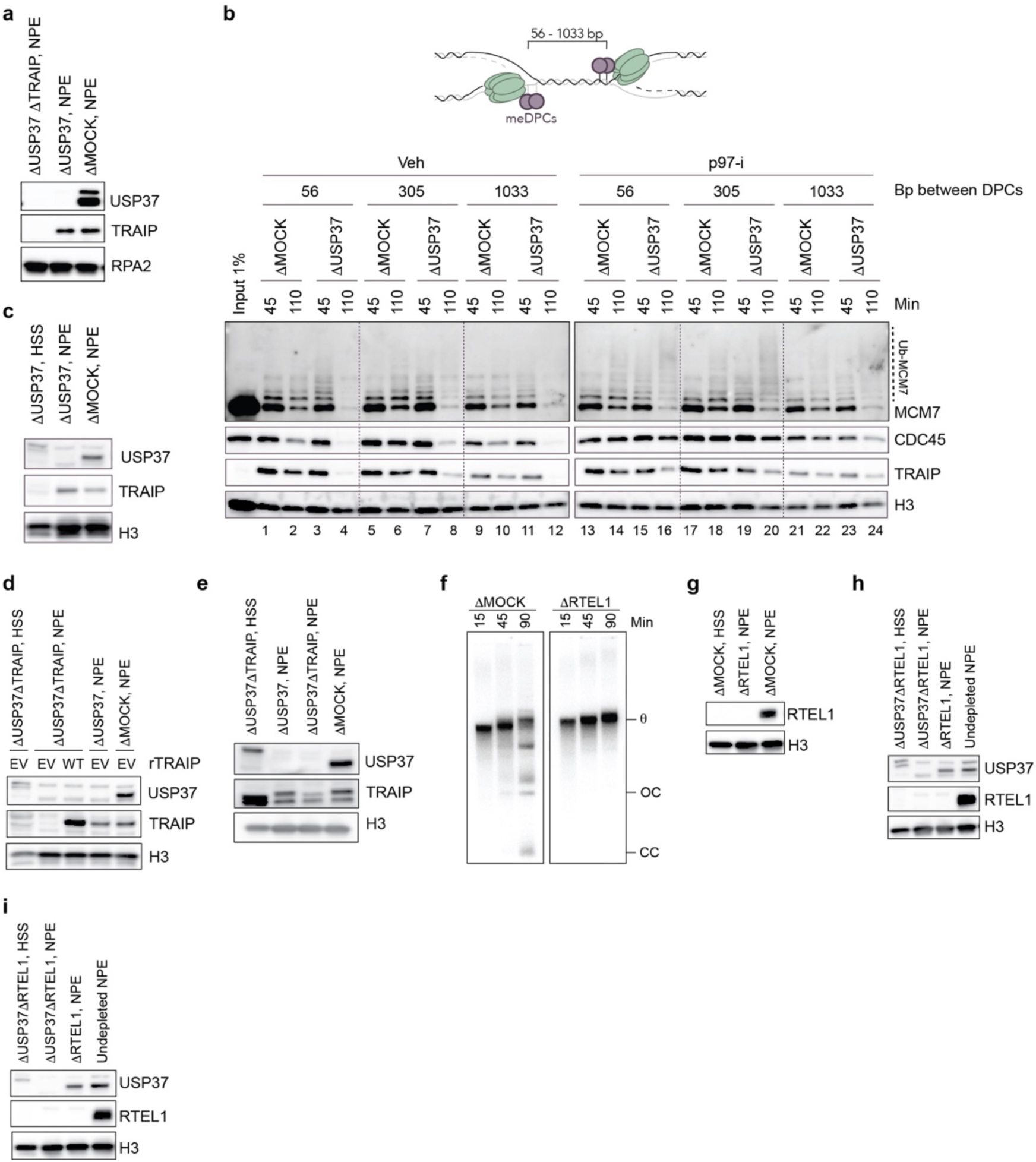
TRAIP promotes premature disassembly of CMGs stalled at distances of up to 1 kb. **a,** Western blot analysis of mock, USP37, and TRAIP depletions. **b,** Top, schematic of the meDPC substrates used, including the variable distance between distal leading strand meDPCs. The indicated meDPC substrates were incubated in egg extracts in the presence or absence of 200 μM p97-i. At the specified times, chromatin was recovered and blotted for the indicated proteins. USP37 depletion efficiency is shown in Extended data Fig. 6c. **c,** Western blot analysis of mock and USP37 depletions. **d,** Western blot analysis of mock, USP37, and TRAIP depletions supplemented with recombinant TRAIP expressed in wheat germ extract. Abbreviations as in Fig. 4b. **e,** Western blot analysis of mock, USP37, and TRAIP depletions. **f,** Plasmid DNA was pre-incubated with LacR and then replicated in the presence or absence of RTEL1 in extracts containing [α-^32^P]dATP. Replication intermediates were separated on a native agarose gel and visualized by autoradiography. RTEL1 depletion delayes progression of replication forks through the LacR array, as seen by stabilization of the theta structure (θ) and absence of fully replicated closed and open circular products (“CC” and “OC”, respectively). The images are part of the same agarose gel, which was cropped to remove irrelevant information. RTEL1 depletion efficiency is shown in Extended data Fig. 6g. **g,** Western blot analysis of mock and RTEL1 depletions. **h,** Western blot analysis of USP37 and RTEL1 depletions. **i,** Western blot analysis of USP37 and RTEL1 depletions.

**Extended data Fig. 7:**
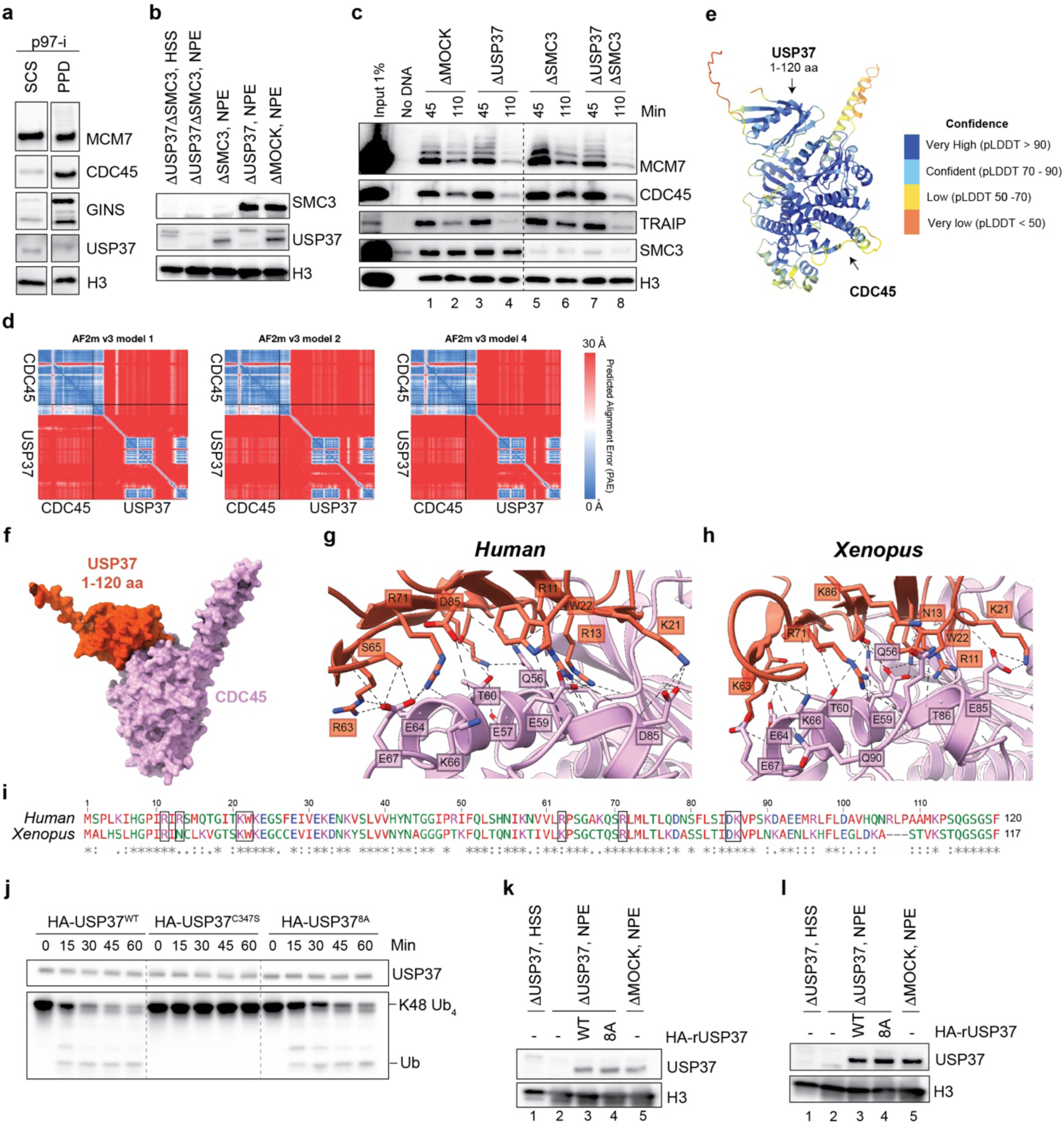
USP37’s PH domain is predicted to interact with CDC45. **a,** Side-by side comparison of USP37 recovery in sperm chromatin spindown (SCS) and plasmid pulldown (PPD) procedures. *Xenopus* sperm chromatin or plasmid DNA were incubated in egg extracts supplemented with indicated inhibitors for 15 min, then recovered and blotted for the indicated proteins. p97-i, NMS-873. The images are all part of the same Western blot, which was cropped to remove irrelevant information. **b,** Western blot analysis of mock-, USP37-, SMC3-, and USP37/SMC3-depletions. Related to Extended data Fig. 7c. **c,** The meDPCs substrate (pDPC4-1033) was incubated in indicated egg extracts. At specified times, chromatin was recovered and immunoblotted for the indicated proteins. USP37 and TRAIP depletion efficiencies are shown in Extended data Fig. 7b. **d,** Predicted alignment error (PAE) plots generated by the 3 AF-M models for the complex of human USP37 and CDC45. **e,** AlphaFold-Multimer (AF-M) prediction of human CDC45 and USP37 interaction colored by pLDDT value, a measure of the confidence of local amino acid positioning. For simplicity, only amino acid residues (aa) 1-120 corresponding to the USP37 PH domain are shown. **f,** Space filling representation of the same model shown in Extended data Fig. 7b colored by chain. **g,** Close-up views of key *Human* USP37 and CDC45 residues that were predicted to interact by AF-M. **h,** Close-up views of key *Xenopus* USP37 and CDC45 residues predicted to interact. **i,** Alignment of human (residues 1-120) and frog (residues 1-117) USP37. Black boxes indicate residues that were substituted to alanines in the USP37^8A^ mutant. Although the residues D85 was not predicted to interact by AF-M v2.3 (panels **d** and **e**), they were predicted to interact by AF-M v2.2 (not shown) and were therefore also mutated. “*”, conserved residues; “:”, conservative substitution; “.”, semi-conservative substitution; “ “, non-conservative substitution; “-“, gap. **j,** A representative in vitro deubiquitylation assay with recombinant *Xenopus* HA-USP37 variants expressed in wheat germ extract. HA-USP37 was immobilized on anti-HA magnetic beads and incubated with K48-linked tetraubiquitin (Ub4). At the indicated times, reactions were stopped with 2x Laemmli sample buffer and blotted for the indicated proteins. **k,** Western blot analysis of mock and USP37 depletions supplemented with recombinant USP37 expressed in wheat germ extract. Abbreviations as in Fig. 5c. **l,** Western blot analysis of mock- and USP37-depletions supplemented with recombinant USP37 expressed in wheat germ extract.

**Extended data Fig. 8:**
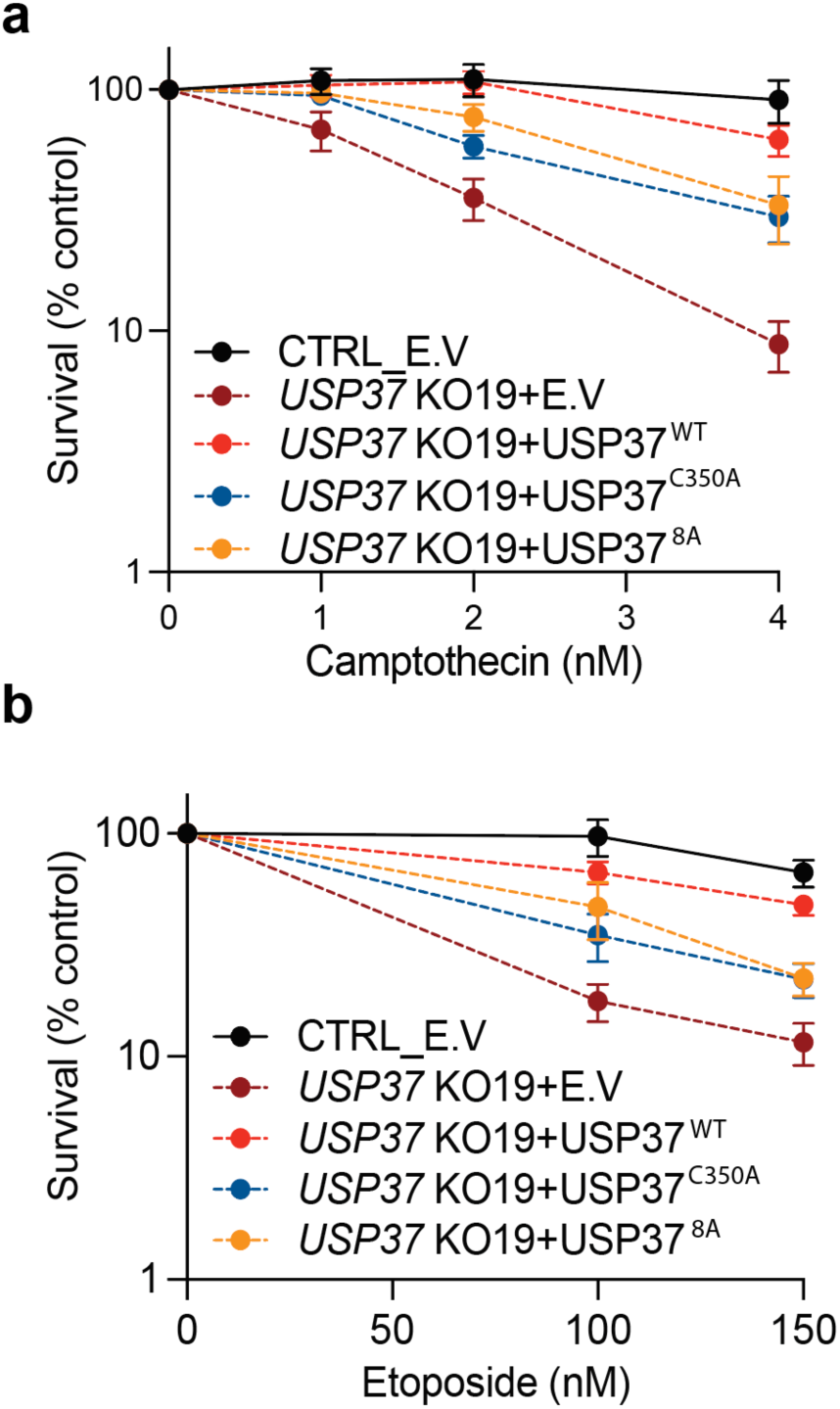
USP37 interaction with CDC45 is required for its protective function towards topoisomerase inhibitors. **a-b,** Clonogenic survival assays of control (CTRL) cells or *USP37* KO19 cells complemented with vectors expressing mCherry (EV), mCherry-USP37^WT^, mCherry-USP37^C350A^ (catalytically inactive), or mCherry-USP37^8A^ (defective for CDC45 interaction) upon treatment with camptothecin (**a**) or etoposide (**b**); n=3 independent experiments. Bars represent means ± SEM.

## Methods

### Animal ethics

Egg extracts were prepared using adult female *Xenopus laevis* (Nasco Cat #LM0053MX). All experiments involving animals were approved by the Harvard Medical Area Standing Committee on Animals (HMA IACUC Study ID IS00000051-6, approved 10/23/2020, and IS00000051-9, approved 10/23/2023). The Harvard Medical School has an approved Animal Welfare Assurance (D16-00270) from the NIH Office of Laboratory Animal Welfare.

### Preparation of DNA constructs

DNA replication experiments in Fig. 3, 5d, and Extended Data Fig. 5 were performed using pKV45 plasmid^56^. DNA replication experiments in Fig. 4d, e and Extended Data Fig. 6f, were performed using pJD156 plasmid^47^. To generate a panel of pmeDPC plasmids, we first generated new variants of pJLS3 plasmid^46^. To this end, we first removed 48xlacO array by digesting plasmid with BsrGI-HF (NEB) and PstI-HF (NEB), followed by blunting and ligation of the digested plasmid using Quick Blunting and Quick Ligation kits (NEB). For pJLS3-56, the resulted pJLS3-no_lacO was first digested with BamHI-HF (NEB) and ApoI-HF (NEB), blunted and ligated using Quick Blunting and Quick Ligation kits (NEB), respectively. The resulted pJLS3-61 was then mutagenized to substitute two nt.BbvCI nicking sites with nt.BspQI sites and to reduce the distance between M.HpaII sites using OK109/OK110 primers (Supplementary Table 2) and a Q5 Site-Directed Mutagenesis Kit (NEB). To generate pJLS3-305, a gene block containing one copy of human ubiquitin gene was introduced into a BssHI/ApoI-digested pJLS3-no_lacO using NEBuilder HiFi DNA Assembly Cloning Kit (NEB). The pJLS3-1033 plasmid was generated by insertion of three additional copies of the human ubiquitin gene into pJLS3-305 in a stepwise manner. pJLS3-305 was first PCR amplified using OK113/OK114 primers (Supplementary Table 2), the second copy of the human ubiquitin gene with a 3’ 15-nt spacer (complementary to OK115 primer) was then introduced into the resulted PCR fragment using NEBuilder HiFi DNA Assembly Cloning Kit (NEB). The resulted pJLS3-562 was first PCR amplified using OK113/OK115 primers (Supplementary Table 2), the third copy of the human ubiquitin gene with a 3’ 15-nt spacer (complementary to OK117 primer) was then introduced into the resulted PCR fragment using NEBuilder HiFi DNA Assembly Cloning Kit (NEB). To generate pJLS3-1033, the resulted pJLS3-805 was first PCR amplified using OK113/OK117 primers (Supplementary Table 2), and the fourth copy of the human ubiquitin gene was cloned into the resulted backbone using NEBuilder HiFi DNA Assembly Cloning Kit (NEB). To generate the pmeDPC-56 substrate, plasmid was first nicked with nt.BbvCI (NEB) and ligated with “Dual-Top/Top-nt.BbvCI” oligonucleotide containing fluorinated cytosines (5’-TCAGCATC[C5-fluor dC]GGTAGCTACTCAATC[C5-fluor dC]GGTACC-3’). The resulted plasmids were then nicked with nt.BspQI (NEB) and ligated with “Dual-Top/Top-nt.BspQI” oligonucleotide containing fluorinated cytosines (5’-CAGCATC[C5-fluor dC]GGTAGCTACTCAATC[C5-fluor dC]GGCTCTTCA-3’) and subsequently crosslinked to methylated M.HpaII-His_6_ as described previously^57^. All other substrates were generated as described above, but both strands were nicked with nt.BbvCI (NEB) and ligated with “Dual-Top/Top-nt.BbvCI” oligonucleotide containing fluorinated cytosines (5’-TCAGCATC[C5-fluor dC]GGTAGCTACTCAATC[C5-fluor dC]GGTACC-3’).

### DNA replication using egg extracts

*Xenopus* egg extracts and de-membranated sperm chromatin were prepared as described^58^. To carry out DNA replication in egg extracts, licensing was first performed by supplementing a high-speed supernatant (HSS) of egg cytoplasmic fraction with plasmid DNA at a final concentration of 7.5 ng/µL (for nascent strand analysis), 15 ng/µL (plasmid pull-down assay) or sperm chromatin at a final concentration of 10,000 sperm/ µL. For replication of LacR-bound plasmids, pJD156 plasmid DNA (300 ng/µL) was mixed with the equal volume of 60 µM LacR^47^ and incubated for 60 minutes at room temperature prior to addition into HSS. Licensing was allowed to proceed for 30 min at room temperature. To inhibit licensing, HSS was pre-incubated for 10 min at room temperature with Geminin, at a final concentration of 10 µM. To initiate replication, 2 volumes of 50% nucleoplasmic extract (NPE) diluted with 1xELB-sucrose (10 mM Hepes-KOH pH 7.7, 2.5 mM MgCl_2_, 50 mM KCl, 250 mM sucrose) was added to 1 volume of licensing reaction. To inhibit CMG assembly, NPE was supplemented with 50 μg/mL recombinant GST-p27^Kip^ (“CDK-i”) or 50 μM PHA-767491 (Sigma-Aldrich PZ0178, “DDK-i”) and pre-incubated at room temperature for 15 mins prior to addition to licensing reaction. To inhibit TOPIIα, replication reactions were supplemented with ICRF-193 (Sigma Aldrich, I4659-1MG) to a final concentration of 200 μM at 5 mins after NPE addition. To prevent nascent strand synthesis in Extended Data Fig. 5d, replication reactions were supplemented with 50 ng/µL of aphidicolin (Sigma Aldrich) at 5 min after NPE addition. For NMS-873 (“p97-i”; Sigma Aldrich, SML1128-5MG) treatment, replication reactions were supplemented with NMS-873 to a final concentration of 200 µM at 40 mins after NPE addition or it was added 5 min prior to replication initiation into NPE (Fig. 4c). For MLN-4924 (“Cul-i”; Active Biochem, A-1139) treatment, NPE was supplemented with MLN-4924 to a final concentration of 400 µM in the replication reaction 5 min prior to replication initiation (Fig. 4c). In Fig. 4d-E, replication reactions where indicated were supplemented with Cyclin B-Cdk1 (Sigma-Aldrich, 14-450M) to a final concentration of 50 ng/μl at 30 min.

For nascent strand analysis in Extended Data Fig. 5a and 6f, replication reactions were supplemented with 0.16 mCi/mL of [a-32P]dATP (Revvity, BLU512H500UC). At the indicated points, samples of the replication reactions were quenched in five volumes of replication stop buffer (80 mM Tris-HCl pH 8.0, 8 mM EDTA, 0.13% phosphoric acid, 10% Ficoll 400, 5% SDS, 0.2% bromophenol blue). Proteins in each sample were digested with 20 mg of Proteinase K (Roche 3115879001) for 1.5 hours at 37°C, and the samples were then resolved on the native 0.8% agarose gel. The dried gels were imaged on the Typhoon FLA 700 PhosphorImager (GE Healthcare).

### Immunodepletions and rescue experiments in egg extracts

For all immunodepletions, 0.5 volumes of the 1 mg/mL affinity-purified antibodies were pre-bound to 1 volume of Dynabeads Protein A (Invitrogen 10002D) by gently rotating at 4°C overnight. 1.5 volumes of undiluted HSS or 50% NPE diluted with 1x ELB were immunodepleted by three rounds of 1-hr incubation with 1 volume of antibody-bound Dynabeads at 4°C. For rescue experiments, one volume of immunodepleted NPE was supplemented with 0.05 volume of wheat germ extract expressing HA-USP37, TRAIP, or wheat germ extract containing an empty vector.

### SDS-PAGE analysis and western blotting in egg extracts

Protein samples were diluted with SDS Sample Buffer to a final concentration of 50 mM Tris [pH 6.8], 2% SDS, 0.1% Bromophenol blue, 10% glycerol, and 5% β-mercaptoethanol and resolved on Mini-PROTEAN or CRITERION precast gels (Bio-Rad) or home-made 6% polyacrylamide gels (MCM7, CDC45). Gels were then transferred to PVDF membranes (Thermo Scientific, PI88518). Membranes were blocked in 5% nonfat milk in 1x PBST for 1 hr at room temperature, then washed three times with 1x PBST, and incubated with primary antibodies diluted to 1:300 – 1:12,000 in 1x PBST containing 1% BSA overnight at 4°C.

Following washes with 1x PBST, membranes were incubated for 1 hr at room temperature with goat anti-rabbit horseradish peroxidase-conjugated antibodies (Jackson ImmunoResearch) at 1:10,000 – 1:30,000 dilution, light chain specific mouse anti-rabbit antibodies (Jackson ImmunoResearch) at 1:10,000 dilution, or rabbit anti-mouse horseradish peroxidase-conjugated antibodies (Jackson ImmunoResearch) at 1:2,000 dilution, diluted in 5% nonfat milk in 1x PBST. Membranes were then washed three times with 1x PBST, developed with SuperSignal West Dura substrate (ThermoFisher), and imaged using an Amersham ImageQuant 800 (Cytiva).

### Antibodies used for western blotting of *Xenopus* proteins

The following rabbit polyclonal antibodies were used for Western blotting: MCM6 (1:5,000; Ref.^7^); MCM7 (1:12,000) and RPA (1:7,000; Ref.^55^; MCM4 (1:2,000; (Bethyl A300-193A)); CDC45 (1:20,000; Ref.^59^); H3 (1:500; (Cell Signaling, 9715S)); TRAIP (1:10,000) and GINS (1:5,000) Ref.^11^; SMC3 (1:5,000; Ref.^60^; SMC2 (1:5000; Ref.^21^); USP37.L (1:5,000; this study); TOP2α (1:5,000; Ref.^47^); RTEL1-N (1:2,500; Ref.^46^). Mouse monoclonal antibodies against the following proteins were used for western blotting: ubiquitin (Santa Cruz Biotechnology, sc-8017 P4D1). Rabbit polyclonal antibodies raised against C terminus of *Xenopus laevis* Usp37.L (Ac-CSQPVSTELNWPTRPPL-OH) were prepared by BioSynth.

### Plasmid pulldown from egg extracts

Plasmid pull-down was performed as described previously^48^. Briefly, 4µl of streptavidin-coated magnetic beads (Dynabeads M-280, Invitrogen) were incubated with 8pmol of biotinylated LacR in 6 volumes of LacR-binding buffer (50mM Tris-HCl [pH 7.5], 150mM NaCl, 1mM EDTA [pH 8.0], 0.02% Tween-20) for 40 min at room temperature. The beads were then washed three times with 5 volumes of 20mM HEPES-KOH [pH 7.7], 100mM KCl, 5mM MgCl_2_, 250mM sucrose, 0.25mg/ml BSA, and 0.02% Tween-20, resuspended in 40µl of the same buffer, and kept on ice. At indicated time points, 6µl of replication reactions were mixed with the prepared beads and rotated for 30 min at 4°C. The beads were then washed three times with 200µl of 20mM HEPES-KOH [pH 7.7], 100mM KCl, 5mM MgCl_2_, 0.25mg/ml BSA, and 0.03% Tween-20. Following complete removal of the washing buffer, beads were mixed with 20µl of 1xSDS Sample buffer and boiled at 95°C for 2 min.

### Sperm chromatin spin-down from egg extracts

Sperm chromatin spin-down was performed as previously described^61^. Briefly, 10 min after replication initiation, 12 μl of replication reaction was mixed with 60 μl of ice-cold 1xELB+ 0.2% TritonX-100, and chromatin and associated proteins were isolated by centrifugation through a 180 μl of sucrose cushion) 1xELB + 0.5M Sucrose). Pelleted chromatin was washed twice with ice-cold 200 μl of 1xELB, resuspended in 2x SDS sample buffer (100 mM Tris pH 6.8, 4% SDS, 0.2% bromophenol blue, 20% glycerol, 10% β-mercaptoethanol). Chromatin and associated proteins were then solubilized by two rounds of sequential boiling at 95°C for 7 min and vortexing for 10 sec prior to western blotting.

### Expression of proteins in wheat germ protein expression system

The wild-type *Xenopus* HA-USP37 open reading frame (ORF) was cloned by introducing the respective gBlocks (IDT) using NEBuilder HiFi DNA Assembly Cloning Kit (NEB) into the AsiSI (NEB) and EcoRI-HF (NEB) double-cut pF3A WG (BYDV) Flexi vector (Promega). The *Xenopus* TRAIP ORF was cloned as above, but the pF3A WG (BYDV) Flexi vector (Promega) was cut with PvuI-HF (NEB) and EcoRI-HF (NEB). The C347S mutation was introduced into the pF3A-HA-USP37 vector using “Round-the-Horn” and primers OK74/OK75 (Supplementary Table 2). To introduce R11A, N13A, K21A, W22A, K63A, R71A, D85A, K86A (8A) mutations into the USP37 gene, pF3A-HA-USP37 vector was first amplified using OK99/OK100 primers (Supplementary Table 2), and the PH-8A gBlock (IDT) (Supplementary Table 2) corresponding to 10 – 285 nt of the USP37 gene (IDT) and containing the indicated mutations was introduced using NEBuilder HiFi DNA Assembly Cloning Kit (NEB). For protein expression, 3 volumes of TnT® SP6 High-Yield Wheat Germ Protein Expression System (Promega) was incubated with 2 volumes of 100 ng/μL purified plasmid at 25°C for 2 hours and used immediately.

### *In vitro* deubiquitylation assay

10 ul of wheat germ reactions expressing HA-USP37 variants were incubated with 25 μl of Pierce™ Anti-HA Magnetic Beads (ThermoFisher) equilibrated into 2xELB-sucrose-BSA-NP-40 buffer (20mM HEPES-KOH [pH 7.7], 100mM KCl, 5mM MgCl_2_, 250mM sucrose) for 1 hr at 4°C, washed five times with 2xELB-sucrose-BSA-NP-40 + 5mM DTT. The beads were then resuspended with 30 μl of 1uM K48-linked tetraubiquitin (Boston Biochem) diluted with 2xELB-sucrose-BSA-NP-40 + 5mM DTT and rotated on the wheel at room temperature. At indicated time points, 5 μl of reactions was mixed with 2x SDS sample buffer (100 mM Tris pH 6.8, 4% SDS, 0.2% bromophenol blue, 20% glycerol, 10% β-mercaptoethanol), and boiled at 95°C for 2 min prior to western blotting.

### AF-M Prediction Generation

Unless stated otherwise all structure predictions were generated using AlphaFold multimer (AF-M)^62^ via ColabFold 1.52 (Ref.^63^). AF-M was run with v2.3 weights1 ensemble, 3 recycles, templates enabled, dropout disabled, and maximum Multiple Sequence Alignments (MSA) depth settings (max_seq = 508, max_extra_seq = 2048). MSAs (paired and unpaired) were fetched from a remote server via the MMSeq2 API^64^ that is integrated into the ColabFold pipeline. To reduce computational burden during the interaction screens we ran 3 out of the 5 AF-M v2.3 models (1,2, and 4) and did not run any pairs where the total amino acid length exceeded 3,600 residues (GPU memory limit).

For Extended Data Fig. 7g-h, five predictions were generated by folding *Human* or *Xenopus* CDC45-USP37 pairs, respectively, using AF-M v2.3 as described above, but with 12 recycles. The top ranked models were then relaxed using AMBER (https://ambermd.org/index.php), and hydrogen atoms were removed with Chimera X1.8. H bonds and salt bridges were visualized with UCSF ChimeraX1.8 using 3Å distance tolerance and 20° angle tolerance criteria. For simplicity, only key residues predicted to interact are shown and/or labeled.

### AF-M Prediction Analysis

Python scripts were utilized to analyze structural predictions generated by AF-M as previously described^50,51^. Briefly, confident interchain residue contacts were extracted from AF-M structures by identifying proximal residues (heavy atom distance <5 Å) where both residues have pLDDT values >50 and PAE score <15 Å. All downstream analysis of interface statistics (average pLDDT, average PAE) were calculated on data from these selected inter-residue pairs (contacts). Average interface pLDDT values above 70 are generally considered confident^65^. The average models score was calculated by averaging the number of independent AF-M models that predicted a specific inter-residue contact across all unique pairs in all models. This number was additionally normalized by dividing by the number of models run to produce a final average model score that ranges from 0 (worst) to 1 (best). An average model value above 0.5 is generally considered confident. pDockQ estimates of interface accuracy scores were calculated independently of the contact analysis described above using code that was adapted from the original implementation as described in Ref.^66^. pDockQ values above 0.23 are considered confident.

### SPOC Interaction Analysis

The random forest classifier algorithm (Structure Prediction and Omics Classifier) SPOC was used to score the binary interaction predicted by via AF-M as described in Ref.^51^. Briefly, this classifier was trained to distinguish biologically relevant interacting protein pairs from non-relevant interaction pairs in AF-M interaction screens. SPOC assigns each interaction a score that ranges from 0 (worst) to 1 (best). Higher scores indicate that AF-M interface metrics and several types of externally omics datasets are consistent with the existence of the binary interaction produced by AF-M. The highest confident interactions are generally associated with SPOC scores above 0.5.

### Cell culture and cell line generation

RPE-1 and HEK293T cells were originally obtained from Prof. Jonathon Pines. RPE-1 *TP53*^KO^ (Ref. ^67^) were cultured in Dulbecco’s Modified Eagle Medium: Nutrient Mixture Ham’s F-12 (DMEM/F-12, Sigma-Aldrich). U2OS Cas9 (Ref.^68^), HEK293T and HEK293T Lenti-X (Takara Bio) cells were cultured in DMEM (PAN-biotech or Sigma-Aldrich). All cell lines were cultured at 37°C and 5% CO_2_ All media were supplemented with 10% (v/v) fetal bovine serum (FBS, BioSera), 100 U/mL penicillin, 100 µg/mL streptomycin (Sigma-Aldrich), 2mM L-glutamine. 10 μg/mL blasticidin (Sigma-Aldrich) was used to select for Cas9 expressing cells. RPE-1 cells stably expressing the mCherry-USP37 vectors were selected with 1 µg/mL G418. USP37 knock-out cells were generated by transient transfection of the sgRNA: AATGTGGTGCTTCGACCCAG targeting exon 5. Three or four days after, single cells were plated, and colonies were picked when visible. Editing was analysed by TIDE (Netherlands Cancer Institute, http://shinyapps.datacurators.nl/tide/) on DNA amplified using the following primers: TGGTCTGTAGTCTAGTCATAGCCT, CCCTTGGTGCAAGATCTCTGT. TRAIP knock-out cells were generated by transduction of the LentiGuide-NLS-EGFP-Puro (kind gift from Durocher laboratory) containing the GACGTGGCCGCCATCCACTG gRNA cloned using the following primers: CACCGACGTGGCCGCCATCCACTG, AAACCAGTGGATGGCGGCCACGTC. Editing was analysed by TIDE on DNA amplified using the following primers: TTGCCCAGGCTAACGGTTTT, AGGCGAAGTATTCACGCTCC.

### CRISPR/Cas9 screens

For CRISPR/Cas9 screens, biological duplicates of U2OS Cas9 wild-type and *USP37* KO cells were transduced at a MOI of 0.3 and 500-fold coverage of the Brunello library^38^. Afterward, transductants were selected with 1.5 µg/mL puromycin for 12 days. Library preparation and next-generation sequencing of the samples was performed as described previously^69^. Guide-enrichment analysis was performed using DrugZ to compare DMSO-treated to etoposide-treated samples^70^.

### Clonogenic survival assays

For clonogenic assays, 500 RPE-1 or U2OS cells were plated in 6 well plates, treated with drugs 24h later and stained and counted 7-10 days later, when visible colonies were formed. Cells were treated with etoposide (VWR International Ltd Cat# CAYM12092-500), camptothecin (C9911, Sigma-Aldrich), talazoparib (Stratech Scientific, S7048-SEL), ICRF-193 (GR332, BIOMOL International), aphidicolin (Merck A0781) or hydroxyurea (H8627, Sigma-Aldrich).

### Competitive cell-growth assay

Control or *USP37* KO human U2OS cells were transfected with sgRNA targeting TRAIP (GACGTGGCCGCCATCCACTG). Samples were collected at the indicated days and subcultured (to prevent confluency) into fresh medium. To calculate the percentage of indels formation, DNA was extracted and regions near the cut-site were PCR-amplified with the indicated primers (Fw: TTGCCCAGGCTAACGGTTTT, Rev: AGGCGAAGTATTCACGCTCC), sequenced, and analyzed by TIDE (Netherlands Cancer Institute, http://shinyapps.datacurators.nl/tide/).

### Plasmid and viral transduction

The plasmids were obtained from VectorBuilder (mCherry, VB230126-1146bke; USP37-WT, VB230119-1435kfc; USP37-C350, VB230126-1139fpu; USP37-8A, VB230119-1445ecf). To generate stable cells, virus was first produced in LentiX 293T cells by co-transfecting the packaging constructs psPAX2 (Addgene #12260) and pMD2.G (Addgene #12259) with the plasmid of interest using TransIT-LT1 (Mirus Bio) according to the manufacturer’s protocol. Viral supernatant was then incubated with cells in the presence of Polybrene (10ug/mL) (Merck), followed by positive selection with Geneticin (Gibco). Transient transfection of Hek293T was obtained using was carried using TransIT-LT1 (Mirus Bio) according to the manufacturer’s protocol. siRNA transfection was performed using Lipofectamine RNAiMAX (Thermo Fisher Scientific) according to manufacturer’s protocol. Cells were seeded for the experiment the following day and were treated and fixed the day after plating.

### Immunoblotting

Cells were lysed in Laemmli buffer (2% sodium dodecyl sulfate (SDS), 10% glycerol, 60 mM Tris-HCl pH 6.8), incubated for 5 min at 95°C and resolved by SDS-PAGE. After transferring onto a nitrocellulose membrane, membranes were blocked with 5% milk or BSA in TBS-T buffer (Tween 20, 0.1%) and incubated with primary antibodies diluted in 5% milk or BSA in TBS-T. Membranes were washed in TBS-T and probed with secondary antibodies. After secondary antibody incubation, membranes were washed in TBS-T, incubated with enhanced chemiluminescence (ECL) mixture for 5 min in the dark, and developed using films or a Chemidoc imaging system (Bio-Rad).

### Immunoprecipitation

For immunoprecipitation, around 2 million Hek293T cells were plated in 10cm dishes. The cells were transfected with the indicated plasmids using MirusLT1 according to manufacturer’s procedure. One days later, cells washed twice with cold PBS and lysed in 1.5mL of IP buffer (20mM Tris HCL pH 7.5, 150mM NaCl, 2mM MgCl_2_, 10% glycerol, 0.5% NP40, and EDTA-free protease and phosphatase inhibitors) and 10uL benzonase (Millipore) for 45 minutes. Lysates were centrifuged at 15,000 rpm for 10 min and supernatants were incubated with 20µL RFP trap magnetic beads (Cromoteck) for 2h at 4C. Samples were washed 4x with IP buffer and finally eluted in 40µL LDS buffer 2x +1mM DTT.

### Immunofluorescence

Cells were plated in 24-well imaging plates and treated with the indicated drugs two days after in the presence of 10uM EdU (A10044, Thermo Fisher Scientific). After treatments, cells were pre-extracted in 0.2% Triton-X in PBS on ice for 5 min and fixed in 4% paraformaldehyde for 10 min. Cells were then blocked in PBS–5% bovine serum albumin (BSA) for at least 30 min. Click reaction was performed to detect S-phase cells with 10 μM AF647-Azide (Jena Bioscience), 4 mM CuSO4 (Sigma-Aldrich), 10 mM (+)-sodium l-ascorbate (Sigma-Aldrich) in a 50 mM Tris buffer. Cells were then incubated with primary antibodies overnight (RPA, NA18 Calbiochem, and γH2AX, 2577 Cell signaling technologies), washed in PBS–0.2% Tween three times and incubated with secondary antibodies for 45 min. After three more washes in PBS–0.2% Tween, nuclei were stained with 4′,6-diamidino-2-phenylindole for 10 min. Images were acquired and analyzed using the Opera Phoenix microscope.

### Proximity ligation assay

The PLA assay was performed as previously described^71^. Briefly, cells were labeled with 10 μM EdU for 10 min, permeabilized 5 min in nuclear extraction buffer on ice (10 mM PIPES, 300 mM sucrose, 20 mM NaCl, 3 mM MgCl_2_, 0.5% Triton X-100) and fixed in 4% PFA for 10 min. When indicated, cells were treated with 1 μM p97i 1 hr and/or with 100 nM CPT 20 min prior to EdU pulse. The click reaction was performed with the Click-iT Kit (Invitrogen C10337, C10339) replacing the supplied Alexa Fluor Azide with a mixture of 20 μM Biotin-Picolyl-Azide (Sigma Aldrich 900912) and 1 μM Alexa Fluor Azide and proceeded according to the manufacturer’s instructions. Cells were blocked in 3% BSA/PBS for 2 hr at room temperature and incubated with primary antibodies overnight at 4°C. For EdU-Flag PLA mouse monoclonal anti-biotin (Jackson Immunoresearch, 200-002-211, 1:2000) and rabbit anti-Flag (Sigma Aldrich F7425, 1:3000) and for EdU-CDC45 PLA mouse monoclonal anti-biotin (Jackson Immunoresearch, 200-002-211, 1:500) and rabbit anti-CDC45 (Cell Signaling Tech #11881, 1:500) were used. After incubation with anti-Mouse MINUS and anti-Rabbit PLUS PLA Probes (Sigma Aldrich DUO82004, DUO82002), cells were subjected to proximity ligation and detection with DUOLINK detection Kit following the manufacturer’s instructions (Sigma Aldrich DUO92008, DUO92014). Coverslips were mounted with Prolong antifade with Dapi (Invitrogen P36935) and cells were imaged on a Nikon Eclipse Ni microscope in conjunction with Elements v4.5 software (Nikon). PLA foci in EdU-positive cells were quantified for each condition and data were analyzed with Prism software.

## Figures and Schematics

Figures were made with Adobe Illustrator. All figures with AF-M predictions were generated in UCSF ChimeraX1.8.

## Statistics and Reproducibility

Graphs and statistical tests were made using Prism v9 (GraphPad Software). Statistics in Fig. 1f-g, 2f-g, 3d, and 5f was calculated using Ordinary one-way-ANOVA. Outliers in Fig. 3d were removed with the ROUT method using Prism. All experiments in *Xenopus* egg extracts were performed at least twice, with a representative result shown.

## Code availability

This paper does not report original code.

