## Supplementary Table 2 for "USP37 prevents premature disassembly of stressed replisomes by TRAIP"

| **Oligonucleotides** | **Source** | **Name** |
| --- | --- | --- |
| AATGTGGTGCTTCGACCCAG | IDT | USP37 sgRNA |
| TGGTCTGTAGTCTAGTCATAGCCT | Sigma | US37 TIDE Fw |
| CCCTTGGTGCAAGATCTCTGT | Sigma | US37 TIDE Rev |
| GACGTGGCCGCCATCCACTG | IDT | TRAIP sgRNA |
| TTGCCCAGGCTAACGGTTTT | Sigma | TRAIP TIDE Fw |
| AGGCGAAGTATTCACGCTCC | Sigma | TRAIP TIDE Rev |
| CTACATGAACGCCATATTGCAATCTC | IDT | OK74 |
| GAGGTGTTTCCGAGGTTACTGAAG | IDT | OK75 |
| TATCGGACCGTGCAGGGAATG | IDT | OK99 |
| GTACCACTGAACAAAGCGGAGAATTTG | IDT | OK100 |
| ACTCAATCCGGCTCTTCAGCCATATGGTGCACTCTCAGTACAATCTGC | IDT | OK109 |
| ACCACCGCGCAAACGCAG | IDT | OK114 |
| TTGTTAGGGAGGAAACCACCGCG | IDT | OK115 |
| TTGGCTTCAACGTAAACCAC | IDT | OK117 |
| CATTCCCTGCACGGTCCGATAGCCATTGCGTGTCTTAAAGTTGGCACTAGTGCGGCAAAGGAGGGTTGCTGTGAGGTGATAGAGAAAGACAATAAATACTCCCTTGTGGTTAACTATAATGCGGGAGGTGGACCAACAAAATTCCAATTGACACAAAACATTAAGACAATTGTGCTGGCGCCTAGTGGCTGCACTCAGTCAGCGTTGATGTTGACTCTGAAGGATGCATCCTCTCTGACTATTGCAGCGGTACCACTGAACAAAGCGGAGAATTTG | IDT | PH-8A gBlock |
| TCAGCATC[C5-fluor dC]GGTAGCTACTCAATC[C5-fluor dC]GGTACC | IDT | Dual-Top/Top-nt.BbvCI |
| CAGCATC[C5-fluor dC]GGTAGCTACTCAATC[C5-fluor dC]GGCTCTTCA | IDT | Dual-Top/Top-nt.BspQI |
| Lincode Non-targeting Control 3 | Dharmacon | Cat# D-001810-03 |
| CAG CAU GGU UAC UAC GAA ATT | Sigma­­­­­^1^ | siTRAIP |
| CGGACCTGTAGCAGTTTCTT | Sigma | TRAIPqPCR Fw |
| CGAAGAAGTCGGAGCAGATAG | Sigma | TRAIPqPCR Fw |
